# Hi-C for genome-wide detection of enhancer-hijacking rearrangements in routine lymphoid cancer biopsies

**DOI:** 10.1101/2025.03.15.643352

**Authors:** Jamin Wu, Shih-Chun A. Chu, Jang Cho, Misha Movahed-Ezazi, Kristyn Galbraith, Camila S. Fang, Yiying Yang, Chanel Schroff, Kristin Sikkink, Michelle Perez-Arreola, Logan Van Meter, Savanna Gemus, Jon-Matthew Belton, Valentina Nardi, Abner Louissant, Dennis Shasha, Aristotelis Tsirigos, Noah Brown, Tatyana Gindin, Marcin P. Cieslik, Minji Kim, Anthony D. Schmitt, Matija Snuderl, Russell J. H. Ryan

**Affiliations:** Department of Pathology, NYU Langone Health, New York, NY; Department of Pathology, University of Michigan Medical School, Ann Arbor, MI; Arima Genomics, Carlsbad, CA; Department of Pathology, Massachusetts General Hospital, Boston, MA; Courant Institute of Mathematical Sciences, Department of Computer Science, New York University, New York, NY; Department of Medicine, Division of Precision Medicine, NYU School of Medicine, New York, NY 10016; Gilbert S. Omenn Department of Computational Medicine and Bioinformatics, University of Michigan, Ann Arbor, MI

## Abstract

Standard techniques for detecting genomic rearrangements in formalin-fixed paraffin-embedded (FFPE) biopsies have important limitations. We performed FFPE-compatible Hi-C on 44 clinical biopsies comprising large B-cell lymphomas (n=18), plasma cell neoplasms (n=14), and other diverse lymphoid cancers, identifying consistent topological differences between malignant B cell and plasma cell states. Hi-C detected expected oncogene rearrangements at high concordance with fluorescence in-situ hybridization (FISH) and supported enhancer-hijacking in recurrent rearrangements of *BCL2*, *CCND1*, and *MYC*, plus unanticipated variants involving homologous loci. Hi-C identified unanticipated non-coding rearrangements involving PD-1 ligand genes and other loci of potential therapeutic relevance, distinguished between functionally divergent classes of *BCL6* rearrangements, and provided topological information supporting the interpretation of atypical *MYC* rearrangements. In biopsies lacking *MYC*-activating rearrangements, Hi-C revealed differential interactions with functionally-validated disease-specific native *MYC* locus enhancers. FFPE-compatible Hi-C detects oncogene rearrangements and their topological consequences at genome-wide scale, finding clinically-relevant drivers that are missed by standard approaches.

## Introduction

The identification of genomic rearrangements is crucial for the precise diagnosis and classification of lymphoid neoplasms^1,2^. Specific oncogene rearrangements define major subtypes of B-cell acute lymphoblastic leukemia / lymphoma (B-ALL)^3–5^ and multiple myeloma (MM)^6,7^ that are used for therapeutic risk stratification. Genomic rearrangements also define distinct subcategories of mature T-cell lymphomas^8,9^ and T-cell acute lymphoblastic leukemia / lymphoma (T-ALL)^10^. In aggressive mature B-cell lymphomas, concurrent rearrangement of the *MYC* and *BCL2* genes identify a high-risk germinal center B cell diffuse large B cell lymphoma (GCB-DLBCL) subgroup^11–14^ that appears to benefit from more intensive chemotherapy regimens^15,16^, while activating *BCL6* rearrangements are characteristic of a genetically and clinically distinctive subgroup of DLBCL^17,18^, and are also associated with poor outcomes when co-occurring with *MYC* rearrangement^14^. Abnormal rearrangements of the immunoglobulin loci commonly result from errors in V(D)J recombination in B cell progenitors or activation-induced cytidine deaminase (AID) activity in germinal center B cells^19,20^, leading to overexpression of partner oncogenes due to the activity of immunoglobulin locus distal enhancers^21^. Similar “enhancer-hijacking” rearrangements are now recognized as recurrent oncogenic events in a range of hematological and solid tumor malignancies involving diverse oncogenes and enhancer-bearing loci^10,22–27^. Unlike gene fusion rearrangements, many enhancer-driven rearrangements do not generate a chimeric transcript that can be identified by RNA-focused methods.

The current clinical standard for detecting genomic rearrangements in routine formalin-fixed paraffin-embedded (FFPE) lymphoma biopsies is fluorescent *in situ* hybridization (FISH). However, FISH is low-throughput and requires the selection of targeted probes, which practically limits its use to testing for a small number of loci in a given disease context. Commonly used FISH probe designs have key limitations, as break-apart probes for a given oncogene locus do not identify the partner locus, while dual fusion strategies often fail to detect variant rearrangements that involve only one of the two targeted loci. Target-capture sequencing of recurrently rearranged noncoding regions has been used to identify rearrangements in FFPE^28,29^, a strategy that can be further enhanced by use of proximity ligation^30^, but such approaches must be tailored to expected rearrangements in a specific disease. Most target-capture and whole-genome sequencing (WGS) strategies rely on short fragments mapping directly to genomic breakpoints for rearrangement detection and thus are prone to artifacts in repetitive intergenic regions, requiring matched normal DNA or other mitigating strategies to minimize such false events ^31,32^. Short-read sequencing also fails to resolve the structure of complex events or confirm cis-interactions between juxtaposed regions. Long-read sequencing technologies and “long-range” techniques such as optical genome mapping (OGM) are not feasible with the fragmented DNA present in FFPE samples.

Hi-C uses proximity ligation to map pairwise topological interactions across the entire genome, revealing topological features such as euchromatin and heterochromatin compartments, topologically associating domains, and selective looping interactions such as those between distal enhancers and promoters ^33,34^. Prior studies using fresh or frozen material have demonstrated the power of Hi-C for genome-wide detection of SVs due to their recognizable effects on spatial DNA proximity ^35–40^, and for detection of gene fusions^41^ and copy number variation^42^ in FFPE tumor samples.

In this study, we performed Hi-C on 44 archival FFPE biopsies of lymphoid cancers. We find that FFPE Hi-C sensitively detects a diverse range of oncogene-activating rearrangements, including clinically significant events that were not identified by routine diagnostic studies, and show that the topological interaction data provided by Hi-C can inform the functional interpretation of complex or uncommon genomic rearrangements.

## Results

### FFPE Hi-C identifies topological features across a range of read depths

We performed Hi-C on FFPE biopsies selected from material at three institutions to compose a diverse cohort of lymphoid cancers with available cytogenetics or FISH data (**Figure 1A**, **Supplementary Tables S1-S2**). Biopsies included mature B and T cell lymphomas that are usually diagnosed via FFPE biopsies, as well as cancers that are more often diagnosed via blood or bone marrow aspirates such as B-ALL, T-ALL and plasma cell neoplasms (PCN), where presentation in a tissue site can present a challenge to standard molecular diagnostic workflows. Our most highly represented tumor subgroups were PCN (n=14; including n=12 cases of MM, and n=2 solitary plasmacytomas that did not progress), and large B-cell lymphoma (n=18; including n=10 systemic GCB-DLBCL, and n=5 primary central nervous system large B cell lymphoma).

**Figure 1.**
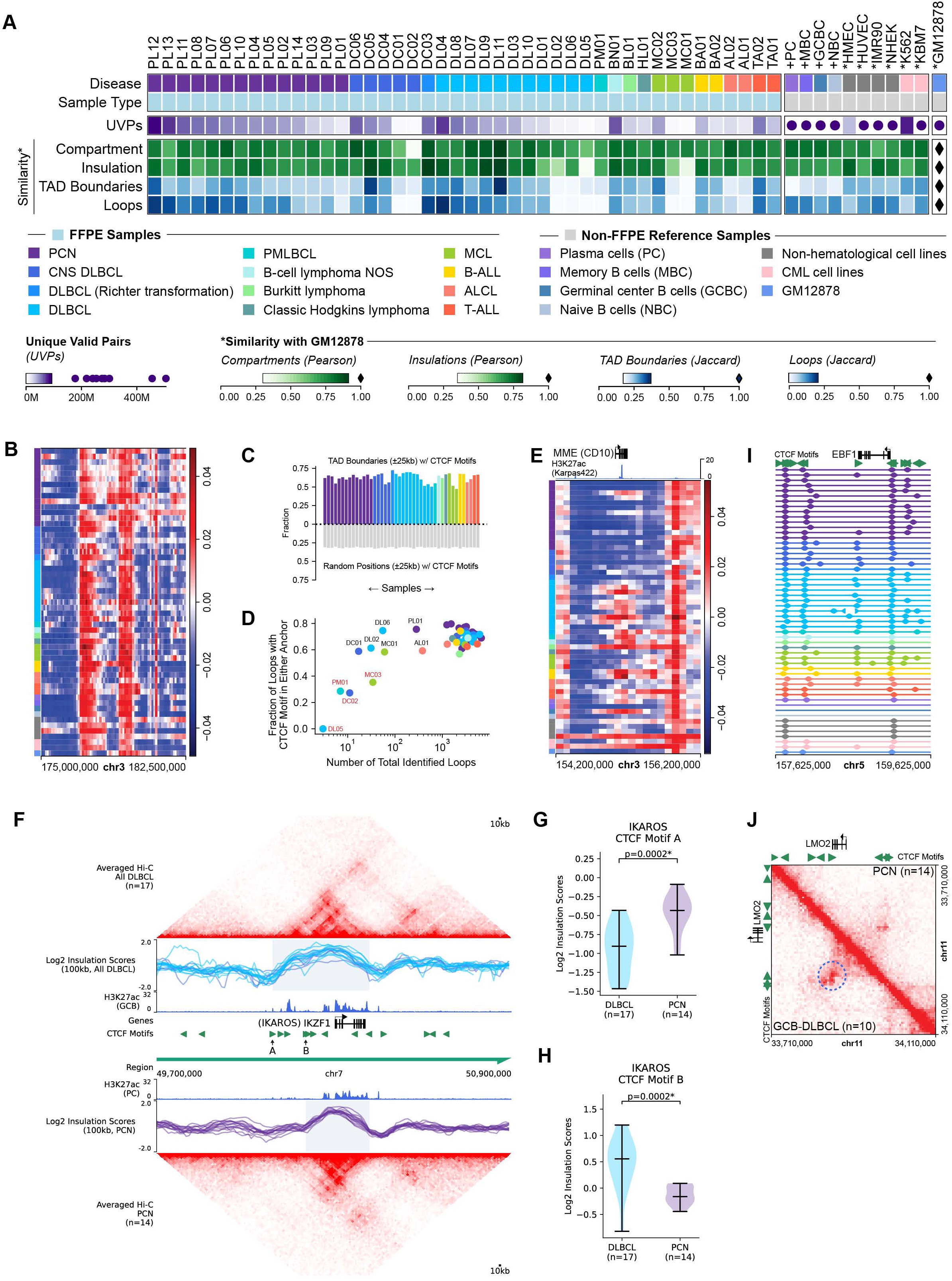
Topological features in FFPE Hi-C datasets. **A,** Overview of sample and reference data characteristics (disease, sample type, number of unique valid read-pairs (UVPs), and similarity metrics compared to non-FFPE reference Hi-C data from the B-lymphoblastoid cell line GM12878. Samples are ordered by diagnostic group (in approximate reverse developmental order for B-cell malignancies) and in descending order of unique valid pairs within each group. Non-FFPE reference samples are prefixed with + (primary samples) or * (cell lines). Legend shows discrete color mappings for diagnostic groups and sample type, and color gradients for each numerical variable as numerical ranges with ● denoting values outside the color range and ⧫ denoting the comparison reference value (GM12878). **B**, Compartment scores in a representative region of chr3 illustrating similarity of A/B compartment scores between samples. **C**, Fraction of TAD boundaries (± one 25kb bin) containing at least one CTCF motif (top) compared with chromosome-matched random positions (bottom). Samples are ordered and colored as in panel A. **D**, Fraction of loops containing at least one CTCF motif in either loop anchor against the number of total identified loops (log scale) in each sample. Samples with <50% of loops containing CTCF motifs in either anchor are labelled in red. Points are colored by diagnostic group as in panel A. **E,** Compartment scores (100kb resolution) demonstrating predominantly inactive/B (blue) compartment states around *MME* (CD10) in PCN and active/A compartment (red) states around *MME* (CD10) in systemic (predominantly GCB type) DLBCL and B-ALL. Samples are ordered and colored as in panel A. **F,** Averaged Hi-C matrix for all DLBCL (top) and PCN (bottom) samples around the *IKZF1* locus, with aligned log2 insulation scores, reference normal population H3K27ac ChIP-Seq read density, and oriented CTCF motifs. CTCF motifs are labeled at local insulation score minima that are specific to DLBCL (“A”) and PCN (“B”). **G** and **H,** Violin plots showing the distribution of Log2 insulation scores in DLBCL and PCN at the two CTCF motifs bordering the region of differential insulation in M. **I**, TAD boundaries at 25kb resolution in the *EBF1* locus, demonstrating a TAD boundary present in DLBCL samples that is absent from most PCN samples. Samples are ordered and colored as in panel A. **J**, Averaged balanced Hi-C matrices at 5kb resolution for GCB-DLBCL (n=10, bottom left triangle) and PCN (n=14, top right triangle) demonstrating a differential loop nearby to LMO2 aligned with two convergent CTCF motifs.

To assess the overall quality of our FFPE Hi-C datasets, we first assessed the similarity of topological features from our cohort with published non-FFPE Hi-C datasets from cell lines and primary B-lineage populations (**Figure 1A**) by correlating all datasets with reference Hi-C from the B-lymphoblastoid cell line, GM12878. The effective sequencing depth of informative read-pairs (unique valid pairs) in our FFPE datasets (median 41.4 million unique valid pairs, range 3.5 – 89.5 million unique valid pairs) was substantially lower than in the non-FFPE reference datasets, a function of both lower raw sequencing depth and a lower proportion of informative reads in the FFPE datasets (**Supplementary Figure S1A and Supplementary Table S3**). Importantly however, we observed that large-scale topological features were well-correlated across our samples and with reference data from non-FFPE samples and cell lines (**Figure 1A** and **Supplementary Figure S1B-D**). A/B compartment scores (100 kb resolution), which represent the broadest measure of nuclear self-association and correspond with packaging of genomic regions into either euchromatin (A compartment) or heterochromatin (B compartment), were well-correlated between samples (**Figure 1A-B**) and robust against low informative read depth (mean Pearson correlation with GM12878 of 0.77, standard deviation 0.09). Insulation scores, which define how frequently adjacent genomic regions interact (100 kb sliding windows), were also well-correlated across the range of sequencing depths in our cohort (mean Pearson correlation 0.69, standard deviation 0.09, **Figure 1A and Supplementary Figure S1B**). Topologically-associating domain (TAD) boundaries and loops are more punctate topological features detectable at higher resolution. Both TAD boundaries and loops were less well correlated in datasets with low effective sequencing depth, and fewer loops were identified in such datasets (**Figure 1A and Supplementary Figure 1E)**. However, all datasets showed enrichment of identified TAD boundaries for motifs of the DNA-binding protein CTCF greater than chromosome-matched random genomic positions (**Figure 1C**) and all but four datasets with low sequencing depth showed the expected strong enrichment of loop anchors for CTCF motifs (including strong enrichment of convergent CTCF motifs for loops containing at least one CTCF motif in each anchor), supporting their biological validity (**Figure 1D** and **Supplementary Figure 1E**). These analyses indicate that topological features shared across different lymphoid cell types are indeed well-preserved and captured by FFPE-compatible Hi-C.

### FFPE Hi-C captures state-specific topological features

Although many genomic regions show similar topological features across different tissues, we expected to see topological differences in some loci that correspond to differences in tissue-specific genome regulatory states. Indeed, principal component analysis of A/B compartment scores showed separation between hematological and non-hematological samples (**Supplementary Figure S1H**). Supervised analysis comparing tumor types identified differential compartment states at loci with known developmental regulation between lymphoid states. For example, *MME* (CD10) showed an active compartment state in B-ALL and germinal center-derived B-cell lymphoma datasets (**Figure 1D**), while *CD22* and *IRF8* showed active compartment states in mature B-cell lymphomas and inactive state in PCN (**Supplementary Figure S1I**), corresponding to expected patterns of developmental and cancer subtype-specific expression of these genes. Conversely, active compartment states were seen in PCN, but not B-cell lymphomas, for genes known to be preferentially expressed in plasma cells such as *HGF* ^43,44^ and *PRDM5* ^45^ (p<0.05 for each gene comparison) (**Supplementary Figure S1J**). Supervised analysis of insulation scores and TAD boundaries highlighted the *IKZF1* locus as a site of differential topology between DLBCL and PCN datasets, with that gene present in a ∼330kb topological domain in DLBCL datasets, but a narrower ∼210 kb topological domain in PCN datasets due to insulation boundary formation at different CTCF motifs in DLBCL versus PCN (**Figure 1F-H**). The genomic region that interacts with *IKZF1* in DLBCL but not PCN contains candidate enhancers that are acetylated in normal germinal center B cells but not plasma cells, suggesting a possible role for this topological boundary shift in state-specific regulation of this key transcription factor, a target of the MM drug lenalidomide^46^.

We also saw significant differences (p<0.05) between PCN and mature B cell neoplasms in the TAD boundaries flanking *EBF1* (**Figure 1I** and **Supplementary Figure S1F** and **S1K**), which encodes a B cell state-defining transcription factor that is downregulated upon plasma cell differentiation ^47^, and a loop specific to GCB-DLBCL biopsies but not PCN (p<0.05) that joined two convergent CTCF motifs around the *LMO2* gene (**Figure 1J** and **Supplementary Figure S1G** and **S1L)**, which encodes a transcriptional regulator that is strongly expressed in germinal center B-cells but not plasma cells ^48^. These findings confirm that regulatory state-specific topological features were well-represented in our FFPE Hi-C datasets, further supporting their quality.

### FFPE Hi-C detects oncogenic structural variants

We next turned our attention to structural variant detection. Hi-C is sensitive for detection of inter-chromosomal and long-range (>100kb) intra-chromosomal SVs because such events result in markedly increased topological interactions between fused genomic regions, accumulating information from read-pairs that map across large regions (**Figure 2A-B**). Overall, we detected SVs across the full spectrum of effective sequencing depths in our cohort with high concordance against findings known from prior clinical testing (**Figure 2A** and **Supplementary Table S2**), including in our shallowest sample with 3.5 million informative read pairs (DL05). Hi-C successfully detected lymphoid cancer subtype-defining gene fusion rearrangements that were expected based on prior clinical assays, such as an *ETV6::RUNX1* fusion in a testicular B-lymphoblastic lymphoma and an *NPM1::ALK* fusion in an ALK+ anaplastic large cell lymphoma (**Figure 2C** and **2D**). Hi-C also detected fusions involving known driver genes that were not identified by prior clinical testing, such as a *DYRK1A::TP63* fusion in a DLBCL biopsy, and an *IGH::RHOH* fusion that linked the 1^st^ intron of the latter gene to the *IGHE* switch region (**Figure 2E** and **2F**).

**Figure 2.**
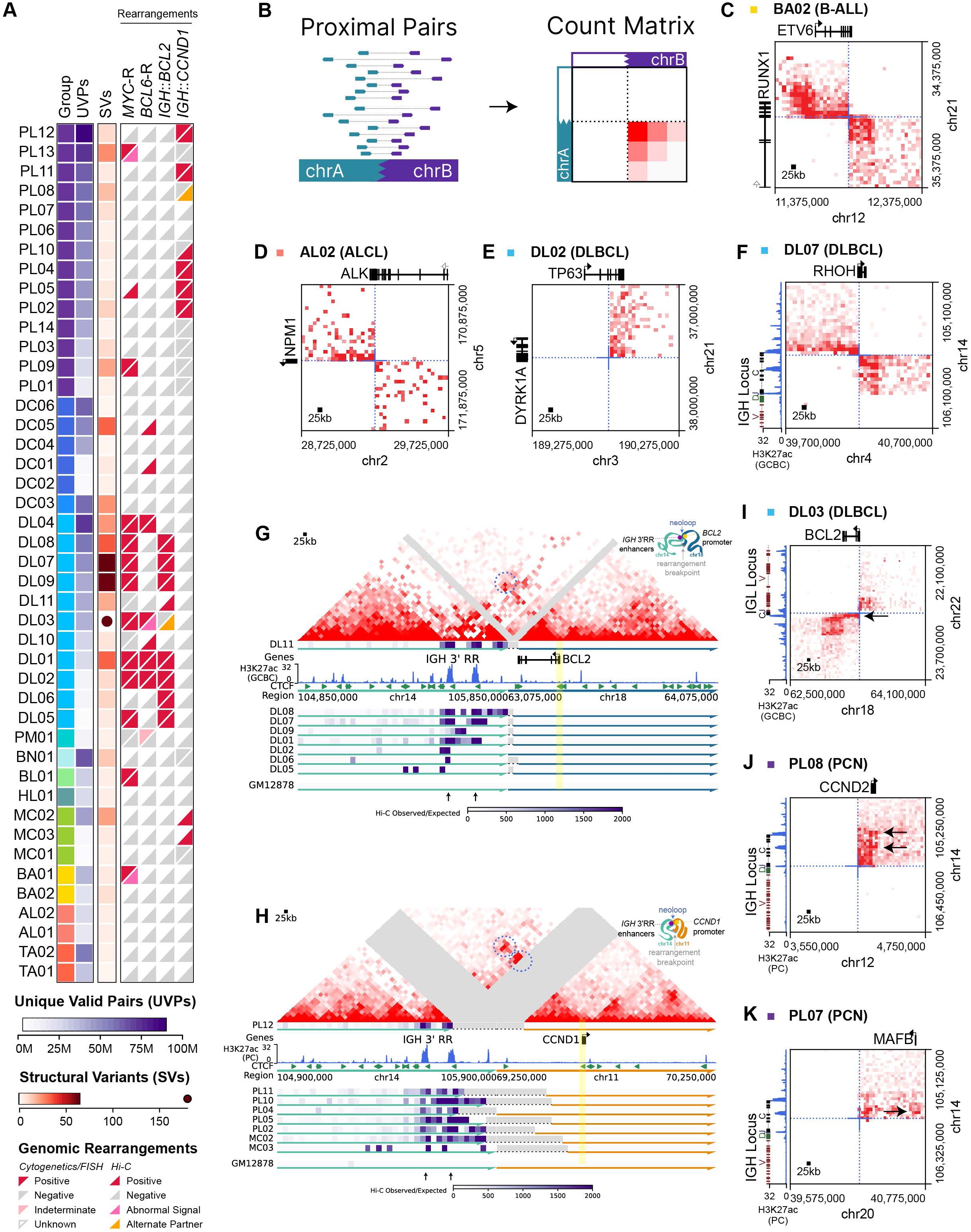
Structural variants detected by FFPE Hi-C in lymphoid biopsies. **A,** Overview of SV detection across the cohort. Disease and UVPs are displayed as in figure 1A for comparison to the total number of SVs and detection of specific rearrangements (*MYC-*R, *BCL6*-R, *IGH::BCL2,* and *IGH::CCND1*) in clinical cytogenetics/FISH testing and from Hi-C. Samples are ordered and colored as in Figure 1A. **B,** Schematic diagram of structural variant detection using Hi-C. **C,D,E,F**, Balanced Hi-C matrices at 25kb resolution showing gene fusions *ETV6::RUNX1* (C), *NPM1::ALK* (D), *DYRK1A::TP63* (E), and *IGH::RHOH* (F) in the indicated biopsies. Reference H3K27ac ChIP-Seq data from normal germinal center B cells (GCBC) is shown for F in the IGH locus. **G**, Top: Balanced Hi-C matrix at 25kb resolution for DLBCL biopsy DL11 showing a reconstructed *IGH::BCL2* rearrangement. A significant neo-loop between an IGH 3’RR enhancer and the *BCL2* promoter is circled. Bottom: virtual 4C tracks (*BCL2* promoter viewpoint, scale is observed / expected Hi-C interactions per 25 kb bin) from 8 DLBCL samples with *IGH::BCL2* rearrangements. Grey regions indicate spacers inserted at breakpoint locations for alignment. A schematic diagram is shown in the upper right corner. **H**, Top: Balanced Hi-C matrix at 25kb resolution for PCN biopsy PL12 showing a reconstructed *IGH::CCND1* rearrangement. Significant neo-loops between *IGH* 3’RR enhancers and the *CCND1* promoter are circled. Bottom: virtual 4C tracks (*CCND1* promoter viewpoint, scale is observed / expected Hi-C interactions per 25 kb bin) for 8 PCN and MCL samples with *IGH::CCND1* rearrangements. Grey regions indicate spacers inserted at breakpoint locations for alignment. A schematic diagram is shown in the upper right corner. **I, J, K,** Balanced Hi-C matrices at 25kb resolution showing putative enhancer-hijacking rearrangements *IGL::BCL2* (I), *IGH::CCND2* (J), and *IGH::MAFB* (K) in the indicated biopsies. Black arrows indicate regions of increased Hi-C interactions between enhancers and oncogene promoters. Reference H3K27ac ChIP-Seq data from normal germinal center B cells (GCBC) or plasma cells (PC) is shown for the IGH and IGL loci.

### FFPE Hi-C supports heterologous IGH enhancer-oncogene interactions

The most frequently recurrent rearrangement pairs in our datasets were *IGH::BCL2*, present in eight cases of GCB-DLBCL, and *IGH::CCND1* present in six MM biopsies and two mantle cell lymphomas (MCL). These well-studied rearrangements rely on long-distance chromatin loops involving the enhancer-donating *IGH* 3’ regulatory regions and / or Mu enhancer (Eμ) to a recipient oncogene to activate its expression ^49^, presenting an opportunity to evaluate the sensitivity of FFPE Hi-C to detect heterologous enhancer-promoter interactions of known oncogenic significance. Indeed, virtual 4C visualization across the breakpoint on the rearranged chromosome pair revealed local peaks of increased topological interactions between the oncogene promoter and known IGH enhancers in all cases (**Figure 2G** and **Figure 2H**), although interactions showed lower signal in cases with lower informative read depth (**Supplementary Figure S2A** and **S2B**). Importantly, Hi-C detected functionally similar variants of these oncogene rearrangements that had not been identified by routine clinical studies. Hi-C identified an *IGL::BCL2* rearrangement in DLBCL biopsy DL03 (**Figure 2I** and **Supplementary Figure S2C**), which was missed by the *IGH::BCL2* dual-fusion FISH strategy used during clinical testing. Hi-C also detected an *IGH::CCND2* rearrangement in MM biopsy PL08 (**Figure 2J** and **Supplementary Figure S2D**), a rare driver event homologous to *IGH::CCND1* that was not evaluated in the FISH panel used clinically for this patient. Both rearrangements showed evidence of new interactions between the oncogene promoter and a partner locus enhancer.

We also saw heterologous *IGH* enhancer interactions associated with *IGH::MAF* and *IGH::MAFB* rearrangements in two MM datasets (**Figure 2K** and **Supplementary Figure S2E, S2F and S2G**). *IGH::MAF* and *IGH::MAFB* rearrangements are both associated with poor prognosis ^7^ and proteosome inhibitor resistance in MM ^50,51^, but many clinical FISH panels do not test for *MAFB* rearrangements due to their rarity, nor the even rarer rearrangements of the homologous gene *MAFA* ^52–54^. For that reason, the *MAFB* rearrangement in case PL07 had not been identified prior to Hi-C.

### FFPE Hi-C identifies enhancer-hijacking rearrangements at diverse loci

While immunoglobulin locus enhancers are known to activate partner oncogenes, systematic identification of enhancer-hijacking rearrangements has been challenging. In addition to identifying genomic fusions, Hi-C may suggest the presence of enhancer hijacking by detection of heterologous interactions between enhancer-rich loci and potential oncogenes ^37,55^. NeoLoopFinder identified 7,864 significant neo-loops across rearrangement breakpoints in the 44 samples in our cohort, which showed punctate interaction signal in aggregate analysis (**Figure 3A**), and were enriched in regions with active enhancer activity based on reference H3K27ac data from germinal center B cells (**Figure 3B**). Neo-loop anchors were enriched in areas joining two active compartments across breakpoints (81.6% of neo-loops), which was not purely attributable to SVs occurring in these regions (56.6% of SVs) (**Figure 3C**).

**Figure 3.**
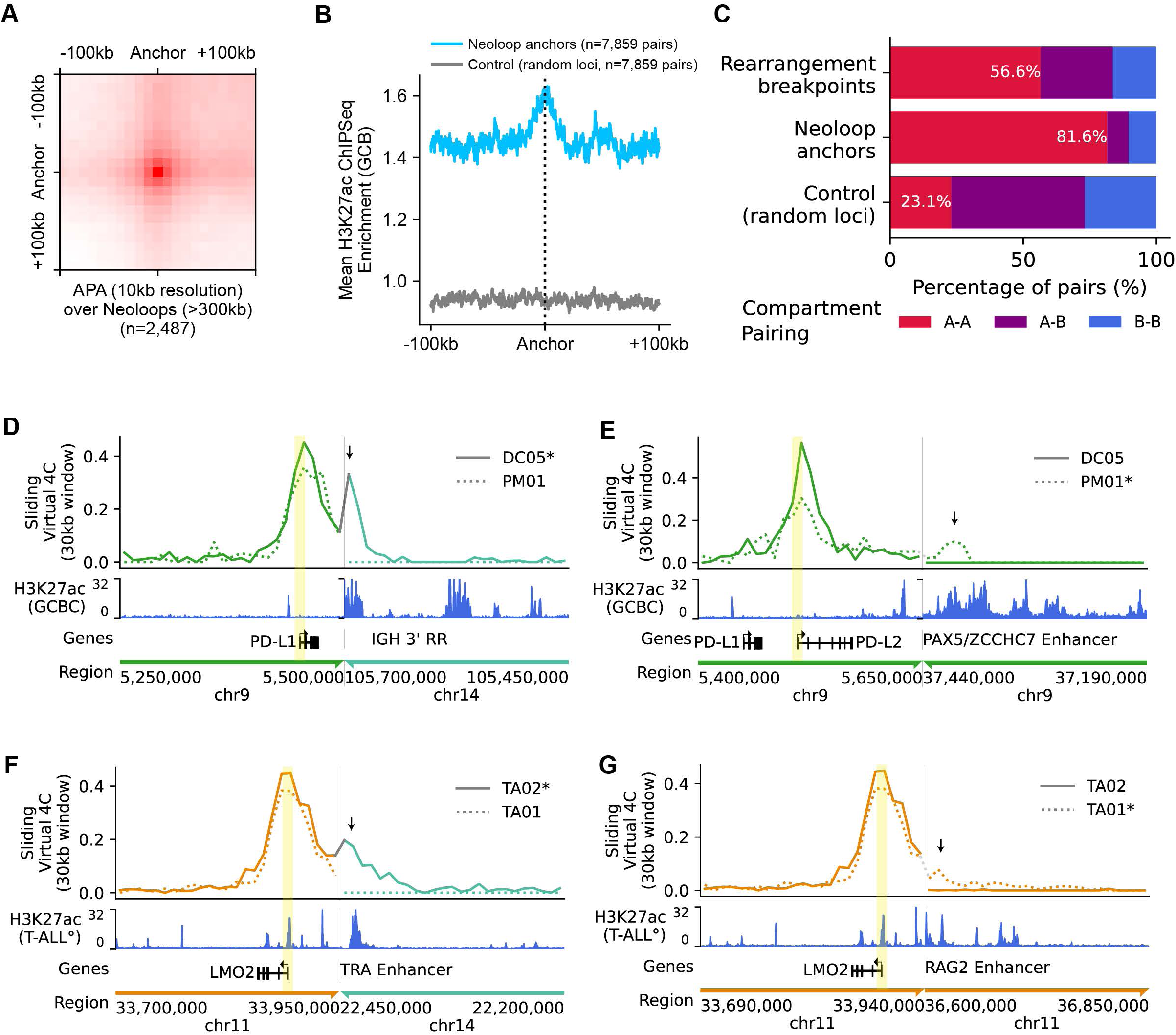
Heterologous interactions across structural variant breakpoints. **A**, Aggregate peak analysis at 10kb resolution with a 100kb radius of neo-loop anchors across all samples with a minimum size of 300kb (n=2,487). **B**, H3K27ac enrichment for reference GCB cells at a 100kb radius around neo-loop anchors across all samples (n=7,859, excluding neo-loop anchors within 100kb of chromosome bounds). **C**, Enrichment of rearrangement breakpoints, neo-loops, and random loci at matched chromosomes for A-A, A-B and B-B compartment pairs. **D,** Virtual 4C analysis across a reconstruction of the chromosomal fusion identified in primary CNS large B cell lymphoma biopsy DC05 between the *CD274* (PD-L1) locus and *IGH*. Hi-C interactions are shown at 10kb resolution using 30kb sliding windows with viewpoint (yellow highlight) at the *CD274* (PD-L1) promoter. Arrow indicates a peak in heterologous interactions with an *IGH* 3’RR enhancer (see reference H3K27ac ChIP-Seq data at bottom). Virtual 4C analysis across the same region in biopsy PM01 (dotted line) is shown for contrast. **E**, Virtual 4C analysis as in D, across a reconstruction of the chromosomal fusion identified in primary mediastinal large B cell lymphoma biopsy PM01 between the *CD274* (PD-L1) / *PDCD1LG2* (PD-L2) locus and *PAX5 / ZCCHC7* locus. Viewpoint (yellow highlight) is at the *PDCD1LG2* (PD-L2) promoter. Arrow indicates a peak in heterologous interactions with *PAX5 / ZCCHC7* enhancers (see reference H3K27ac ChIP-Seq data at bottom). Virtual 4C analysis across the same region in biopsy DC05 (solid line) is shown for contrast. **F,** Virtual 4C analysis across a reconstruction of the chromosomal fusion identified in T-ALL biopsy TA02 between the *LMO2* and *TRA/D* loci. Hi-C interactions are shown at 10kb resolution using 30kb sliding windows with viewpoint (yellow highlight) at the *LMO2* promoter. Arrow indicates a peak in heterologous interactions with an *TRA/D* locus enhancer (see reference T-ALL H3K27ac ChIP-Seq data at bottom). Virtual 4C analysis across the same region in biopsy TA01 (dotted line) is shown for contrast. **G**, Virtual 4C analysis as in F across a reconstruction of the deletion / intrachromosomal fusion identified in T-ALL biopsy TA01 between the *LMO2* and *RAG2* loci. Hi-C interactions are shown at 10kb resolution using 30kb sliding windows with viewpoint (yellow highlight) at the *LMO2* promoter. Arrow indicates a peak in heterologous interactions with a *RAG2* locus enhancer (see reference T-ALL H3K27ac ChIP-Seq data at bottom). Virtual 4C analysis across the same region in biopsy TA02 (solid line) is shown for contrast.

Checkpoint inhibitor therapies targeting programmed death-1 (PD-1) have shown clinical benefits in classic Hodgkin lymphoma ^56^ and primary mediastinal large B-cell lymphoma ^57,58^, which show frequent amplification and genomic rearrangement of the locus encoding the adjacent genes for PD ligand 1 (*CD274*) and PD ligand 2 (*PDCD1LG2*) on chromosome 9p24.1 ^59,60^. These genes undergo activating rearrangement at a lower frequency in other DLBCL subtypes, but clinical testing is not routinely performed. FFPE Hi-C data identified two cases of large B cell lymphoma with genomic rearrangements of 9p24.1, one that juxtaposed *CD274 t*o *IGH* locus enhancers in a primary CNS large B-cell lymphoma, and one linking *PDCD1LG2* to the well-characterized *PAX5/ZCCH7* locus super-enhancer in a primary mediastinal large B-cell lymphoma, a recurrent rearrangement partner for other DLBCL oncogenes such as *MYC* (**Figure 3D** and **3E**, and **Supplementary Figure S3A** and **S3B)**. In both cases, increased topological interactions were seen between the oncogene promoter and partner locus enhancers and immunohistochemistry confirmed expression of PD-L1 in the former case (**Supplementary Figure S3C)**.

T-ALL is a disease with a high frequency of non-coding genetic driver events. FFPE Hi-C analysis of two T-ALL case revealed clinically undetected rearrangements that juxtaposed a known T-ALL oncogene, *LMO2*, to known recurrent partner loci, *TRA* and *RAG2* (**Figure 3F** and **3G, Supplementary Figure S3D** and **S3E**). Both rearrangements showed increased topological interactions between *LMO2* and strong distal enhancers in the partner locus. The presence of an *LMO2* rearrangement is important for classification into recently-defined genomic subtypes of T-ALL^10^. The ∼3 Mb intrachromosomal deletion of chr11 that juxtaposes *RAG2* enhancers to the *LMO2* gene is exclusively seen in the *TAL1* DP-like subtype, which shows strong RAG gene expression, and is associated with worse event-free survival within this group^10^.

### FFPE Hi-C detects large oncogenic copy number alterations

Detection of the *RAG2::LMO2* enhancer-hijacking event in TA01 highlights the ability of Hi-C to identify intragenic SVs. For intragenic SVs that alter chromosomal copy number (deletions and amplifications), Hi-C can detect such events via either genomic fusion or copy number analyses (**Supplementary Figure S3F)**. *LMO2*-activating events in T-ALL usually co-occur with lesions that activate expression of the *TAL1* gene, the most common of which is an 90kb deletion that fuses the adjacent *STIL* and *TAL1* genes. Hi-C signal in this region was suggestive of a *STIL::TAL1* fusion in case TA02 which had been detected by clinically performed genomic microarray, although the size of this event was too small for detection using our 25kb resolution Hi-C-derived copy number analysis (**Supplementary Figure S3G)**. Hi-C-derived copy number profiles in other cases indicated other SVs with known disease associations and potential prognostic significance. Hi-C suggested the presence of *CDKN2A/B* loss in both of our T-ALL samples for which prior DNA microarray testing had revealed *CDKN2A/B* deletions, as well as in three additional samples for which prior clinical testing for *CDKN2A/B* loss had not been performed (one DLBCL, B-ALL, and a B-cell lymphoma not otherwise classifiable) (**Supplementary Figure S3H**). Additionally, five of our DLBCL cases showed SVs resulting in focal copy gain of 2p16.1 containing *REL* **(Supplementary Figure S3I),** a common target of focal genomic amplifications in GCB-DLBCL and PMBL ^61,62^, although the specific oncogenic function of this lesion remains controversial ^63^.

### Hi-C distinguishes *BCL6* gene-activating from *BCL6*-LCR-donating events

Recent studies have found that a subset of oncogene loci can undergo two functionally distinct types of rearrangements, one in which the oncogene is transcriptionally activated by partner locus regulatory elements, and one in which that same oncogene locus donates active enhancers to drive expression of a different oncogene in the partner locus. For example, *BCL11B* can act as either a recipient oncogene ^24^ or enhancer-donating locus ^10,64^ in rearrangements that occur in phenotypically distinct myeloid and / or T-cell progenitor leukemias, while the *MYC* locus, a recipient oncogene in many lymphoid cancers, can act as an enhancer donor in myeloid progenitor-derived leukemias ^24,65^. Similarly, *BCL6* is known to undergo rearrangements that activate *BCL6* gene expression via promoter substitution ^66,67^ and structurally distinct “pseudo-double hit” rearrangements in which the *BCL6* distal super-enhancer / locus control region (LCR), active only in the setting of a strong germinal center gene expression program, is donated to activate MYC^29,68–71^. Standard clinical *BCL6* locus break-apart FISH assays cannot distinguish between these rearrangement types; however, these functionally distinct events were distinguishable in our Hi-C data (**Figure 4A**). Three DLBCL cases showed classic *BCL6* promoter-swap rearrangements that have been implicated in *BCL6* overexpression in non-germinal center type DLBCL, with a genomic fusion linking the *BCL6* first intron to promoter-like switch regions in immunoglobulin loci (*IGH* and *IGL*), or the *JCHAIN* promoter (**Figure 4B, 4C** and **4D** and **Supplementary Figure S4A**, **S4B**, **S4C**). This structure contrasted with rearrangements in two other GCB-DLBCL biopsies^71^ that linked the *BCL6* LCR to the *MYC* gene (**Figure 4E** and **4F** and **Supplementary Figure S4D** and **S4E**), where breakpoints in both loci were intergenic, and new topological interactions were seen between the *BCL6* enhancer and *MYC* promoter.

**Figure 4.**
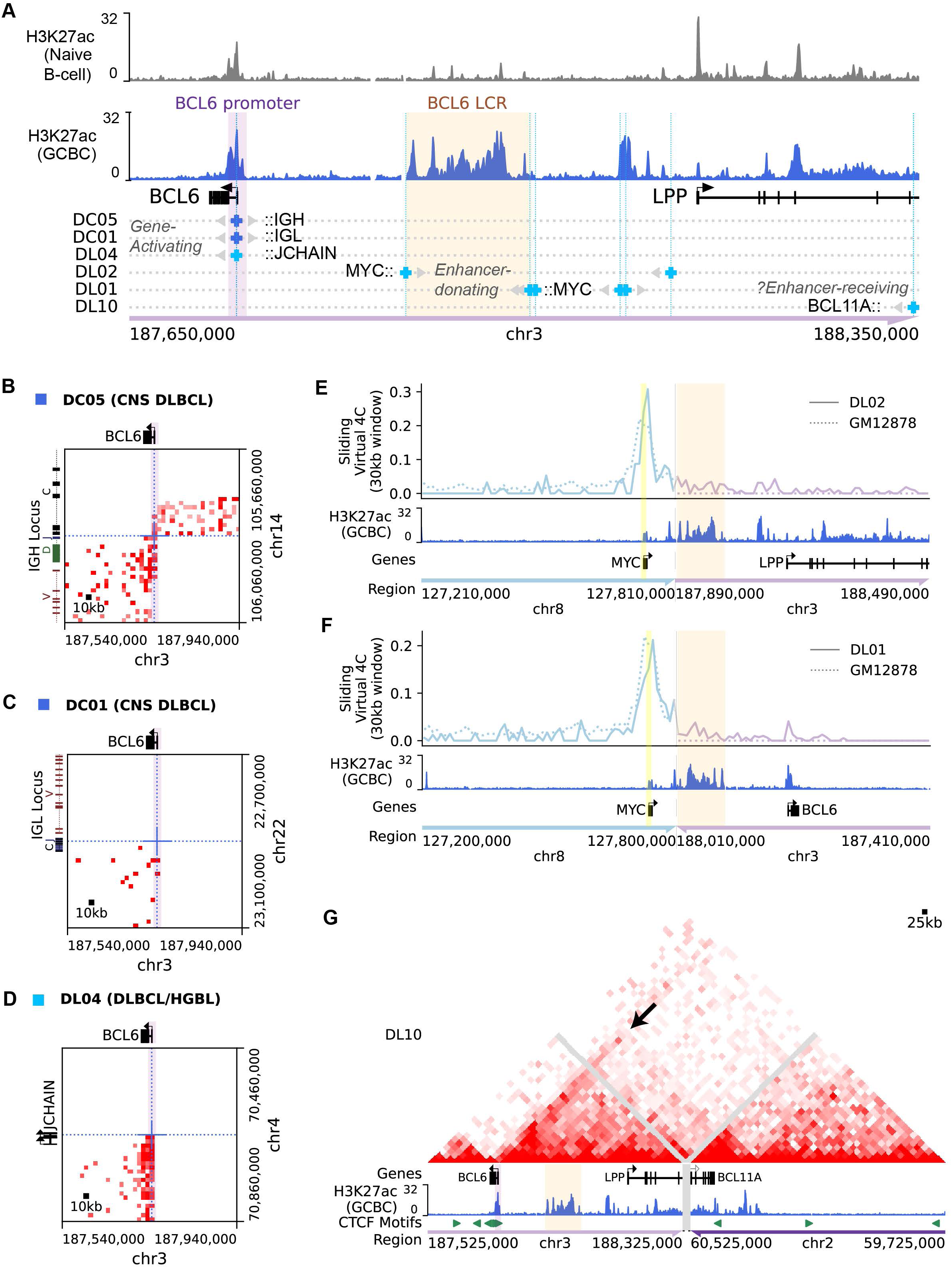
Distinct classes of *BCL6* rearrangement detected by Hi-C. **A**, Location of chromosomal fusion breakpoints in two PCNSL and four systemic DLBCL samples with *BCL6* / *LPP* locus rearrangements. Reference naïve B-cell and GC B cell H3K27ac ChIP-Seq data is shown at top. The locations of the *BCL6* promoter / intronic super-enhancer and distal *BCL6*-LCR super-enhancer are highlighted. At breakpoints, gray arrows indicate one or both genomic regions that are fused to a partner locus across that breakpoint. Genes in the partner locus are indicated in black text. **B, C, D**, Balanced Hi-C matrices at 10kb resolution showing rearrangements between the *BCL6* promoter / first intron and *IGH*, *IGL*, or *JCHAIN* loci in the indicated samples. **E, F**, Virtual 4C analysis across reconstructions of chromosomal fusions identified in two samples with chromosomal fusions that juxtapose *MYC* with the *BCL6*-LCR enhancer complex, DL02 (E) and DL01 (F). Hi-C interactions are shown at 10kb resolution using 30kb sliding windows with viewpoint (yellow highlight) at the *MYC* promoter. Virtual 4C analysis across the same region in GM12878 (dotted line) is shown for contrast. **G**, Balanced Hi-C interaction matrix at 25kb resolution across a reconstruction of the chromosomal fusion between the *BCL6* / *LPP* and *BCL11A* loci in DLBCL biopsy DL10. Reference GC B cell H3K27ac ChIP-Seq data and CTCF motif tracks are shown below. An arrow points to a local peak of interaction between the *BCL6* promoter and a candidate *BCL11A* enhancer present in H3K27ac data.

Hi-C data also revealed two cases with rearrangements of the *BCL6* locus that did not fit either of these functionally characterized patterns. In DL10, the *BCL6* locus was rearranged to an amplified 2p16 locus containing *BCL11A*, *REL*, and *XPO1*, a common site of DLBCL SVs as noted previously. Analysis of the Hi-C interaction matrix revealed new enhancer-promoter interactions between a candidate enhancer adjacent to *BCL11A* and the *BCL6* promoter (**Figure 4G** and **Supplementary Figure S4F**). Although *BCL6*-activating rearrangements are difficult to functionally validate due to strong rearrangement-independent *BCL6* expression in a large proportion of DLCBL^72^, these topological findings suggest that this specific event might contribute to both overexpression of 2p16 genes (due to amplification) and transcriptional dysregulation of *BCL6* (via heterologous enhancer interactions). Notably, a recent large-scale effort to map *MYC* rearrangement partners identified two DLBCLs with rearrangements that appear to juxtapose this same region of the *BCL11A* locus to *MYC* ^29^, further supporting this region as a possible donor of oncogene-activating enhancers. The other event was in case DL03, in which break-apart FISH had reported a *BCL6* rearrangement, but Hi-C revealed the actual lesion to be a complex SV that linked a focal segment of the *LPP* gene to two loci on the short arm of chr3 **(Supplementary Figure S4G and S4H)**. Neither partner locus contains a known oncogene, and there was no evidence of increased topological interactions between the *BCL6* gene and either partner locus, nor significant heterologous interactions of the *BCL6* LCR. Although an oncogenic function of this lesion cannot be ruled out, it seems more likely to be a passenger event, particularly since both proximal and distal portions of the *BCL6* locus are known to be hotspots for off-target DNA damage mediated by activation-induced cytidine deaminase (AID), Thus, Hi-C can effectively distinguish *BCL6* locus rearrangements of known oncogenic function from events of unclear significance, while FISH cannot.

### FFPE Hi-C identifies enhancer-donating partners for *MYC* rearrangements

Activating *MYC* rearrangements are known to be diverse in both structure and partner loci ^28^. *MYC* locus rearrangements were detected in 12 of our biopsies (8 DLBCL, 1 Burkitt lymphoma, 2 MM, and 1 B-ALL). Consistent with prior findings ^28,29^, some *MYC* rearrangement breakpoints clustered adjacent to or within the 5’ end of the *MYC* gene, while others were scattered in intergenic regions up to 1 Mb from *MYC*, primarily on the 3’ side (**Figure 5A** and **Supplementary Figure S5A-G**). The DLBCL cases we evaluated by Hi-C were enriched for known double-hit lymphomas, which more frequently involve non-*IGH MYC* partners, ^29^ allowing us to explore the topology of diverse *MYC* partner loci. Breakpoint detection identified rearrangements between *MYC* and *IGH* in two MM samples, one DLBCL, and one Burkitt lymphoma. Three of these were simple rearrangements with evidence of new interactions between the *MYC* promoter and *IGH* 3’ regulatory region enhancers (**Figure 5B** and **Supplementary Figure S5B**). The fourth case, MM biopsy PL05, showed a complex three-way rearrangement involving the *IGH*, *CCND1*, and *MYC* loci in cis, which could be reconstructed based on the relative strength of topological interactions between all three loci. Hi-C derived copy number amplifications of chr11 and chr14 also supported the reconstructed derivative chromosome (**Supplementary Figure S5H-I**).

**Figure 5.**
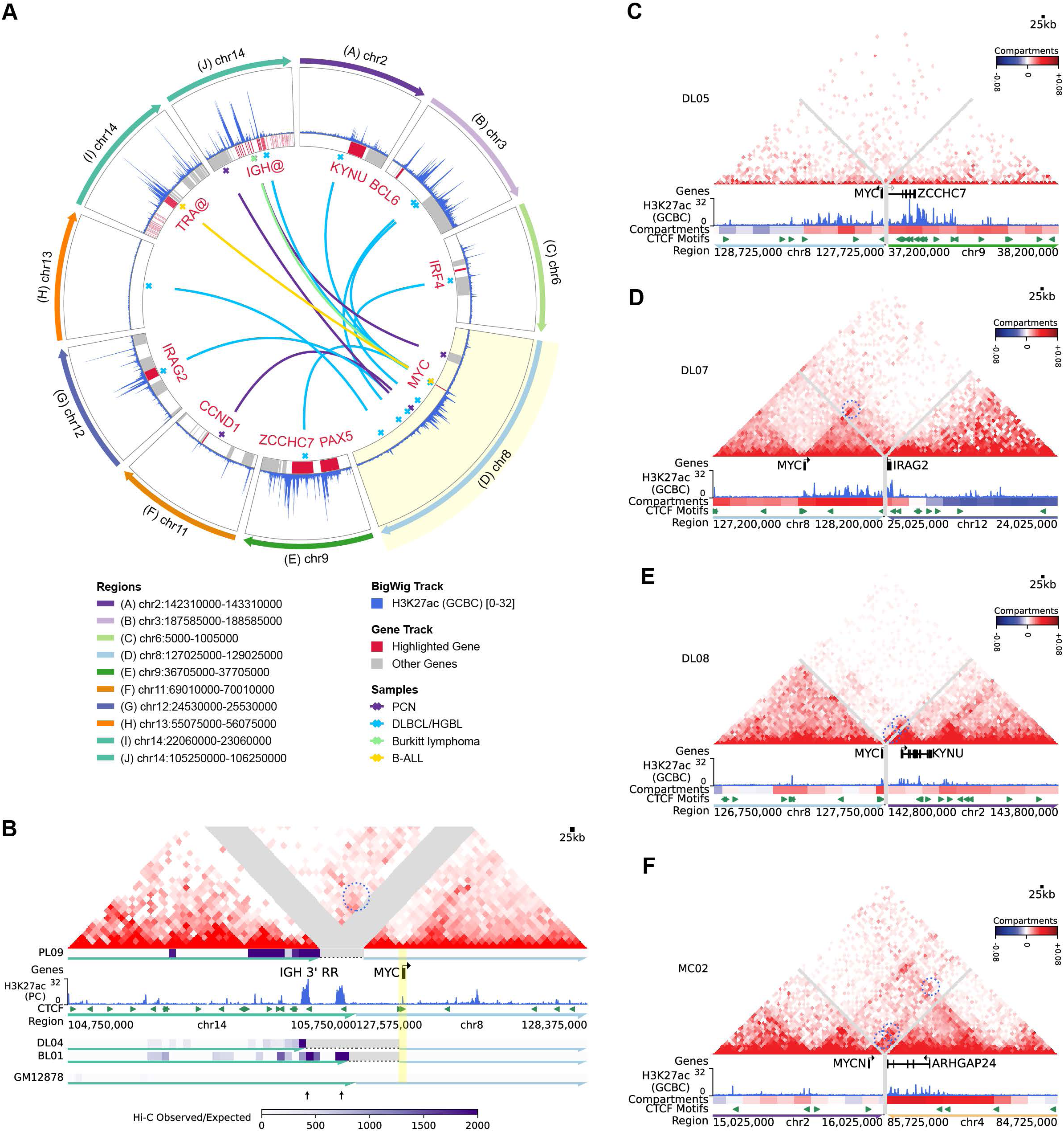
Enhancer-hijacking *MYC* rearrangements detected by Hi-C. **A**, Circos plot depicting inter-chromosomal *MYC* locus rearrangements identified by Hi-C across the cohort. Rearrangements are color-coded by cancer type. Reference H3K27ac data is from normal GC B cells. **B**, Top: Balanced Hi-C matrix at 25kb resolution for PCN biopsy PL09 showing a reconstructed *IGH::MYC* rearrangement. A significant neo-loop between an IGH 3’RR enhancer and the *MYC* promoter is circled. Bottom: aligned virtual 4C heatmaps (*MYC* promoter viewpoint, scale is observed / expected Hi-C interactions per 25 kb bin) from three samples with simple IGH::MYC rearrangements. Grey regions indicate spacers inserted at breakpoint locations for alignment. Reference H3K27ac ChIP-Seq track from normal plasma cells and CTCF motifs are also aligned. **C, D, E,** Balanced Hi-C matrices at 25kb resolution showing reconstructed rearrangements of *MYC* with the *PAX5/ZCCHC7* (C)*, IRAG2* (D) and *KYNU* (E) loci in the indicated DLBCL samples. Blue circles indicate significant neo-loops supportive of enhancer hijacking interactions between *MYC* and partner locus enhancers. Tracks at bottom include Hi-C derived compartment score for the same sample (red = A compartment, blue = B compartment), reference normal GC B cell H3K27ac ChIP-Seq, and CTCF motifs. **F**, Balanced Hi-C matrix at 25kb resolution showing a reconstructed rearrangement between the *MYCN* and *ARHGAP24* loci in mantle cell lymphoma biopsy MC02. Significant neo-loops involving the *MYCN* promoter region are indicated with blue circles. Hi-C compartment scores from the same sample, H3K27ac ChIP-Seq signal, and CTCF motifs are aligned at bottom.

The eight *MYC* rearrangements to non-immunoglobulin loci included rearrangements with well-described recurrent partner enhancers, including the previously mentioned *BCL6* locus control region (n=2) and the *PAX5* / *ZCCH7* super-enhancer (**Figure 5C**). Two other *MYC* partners identified by Hi-C in DLBCL biopsies that have been identified in prior studies, the *IRAG2* ^29^ and *KYNU* ^30^ loci, showed strong candidate enhancers (based on H3K27ac ChIP-Seq data from normal germinal center B cells), an active “A” compartment state, and significant neo-loops linking the candidate enhancers with the *MYC* promoter, suggesting that these are *bona fide* enhancer-donating loci (**Figure 5D** and **Figure 5E**). In contrast, Hi-C did not clearly resolve enhancer-promoter interactions for a rearrangement between *MYC* and the *TRA* locus, a recurrent rearrangement partner, in B-ALL lymph node biopsy BA01 (**Supplementary Figure S5F**). This was likely in part due to the sub-clonal nature of the rearrangement in this sample (approximately 15% of tumor cells by FISH) as further detailed in the discussion (**Supplementary Figure S5J)**.

Interestingly, blastoid mantle cell lymphoma biopsy MC02 showed a rearrangement involving the locus of the *MYC* homolog *MYCN* and a candidate enhancer adjacent to the *ARHGAP24* locus (**Figure 5F** and **Supplementary Figure S5G**), which was previously identified as a *MYC* rearrangement partner in a case of mantle cell lymphoma ^73^. This case displayed the formation of a strong neo-TAD anchored by convergent CTCF loops across the rearrangement breakpoint, constraining multiple significant neo-loop interactions between the *MYCN* promoter and the candidate enhancers in an active “A” compartment state. Together, the *MYC* and *MYCN* rearrangement events seen in our Hi-C cases illustrate how Hi-C-derived topology can support the function of uncommon enhancer-hijacking rearrangements.

In contrast to the cases discussed above, DLBCL cases DL03 and DL09 showed rearrangements involving the *MYC* locus that lacked clear evidence for enhancer hijacking. In DL09, a breakpoint approximately 860kb downstream of *MYC* was fused to a region of chr13 with a heterochromatic B compartment state that entirely lacked candidate enhancers in reference GCB and Karpas-422 H3K27ac ChIP-Seq data (**Figure 6A** and **Supplementary Figure S6A**). DL03 showed a complex *MYC* rearrangement in the context of chromosome 8 chromothripsis (**Figure 6B and Supplementary Figure S6B-D**), with a partner locus that showed minimal evidence of enhancers in reference GCB and Karpas 422 H3K27ac data, nor topological looping to the *MYC* promoter. To our knowledge, the rearrangement partner loci in these cases have not been previously identified as oncogene rearrangement partners, including in a recent large-scale rearrangement mapping effort in DLBCL ^29^. Both cases showed a large number of genome-wide SVs, with chr8 chromothripsis present in DL03 and chr20 chromothripsis in DL09 (**Supplementary Figure S6D-E)**, suggesting that these *MYC* locus SVs might represent passenger events rather than functional *MYC*-activating rearrangements.

**Figure 6.**
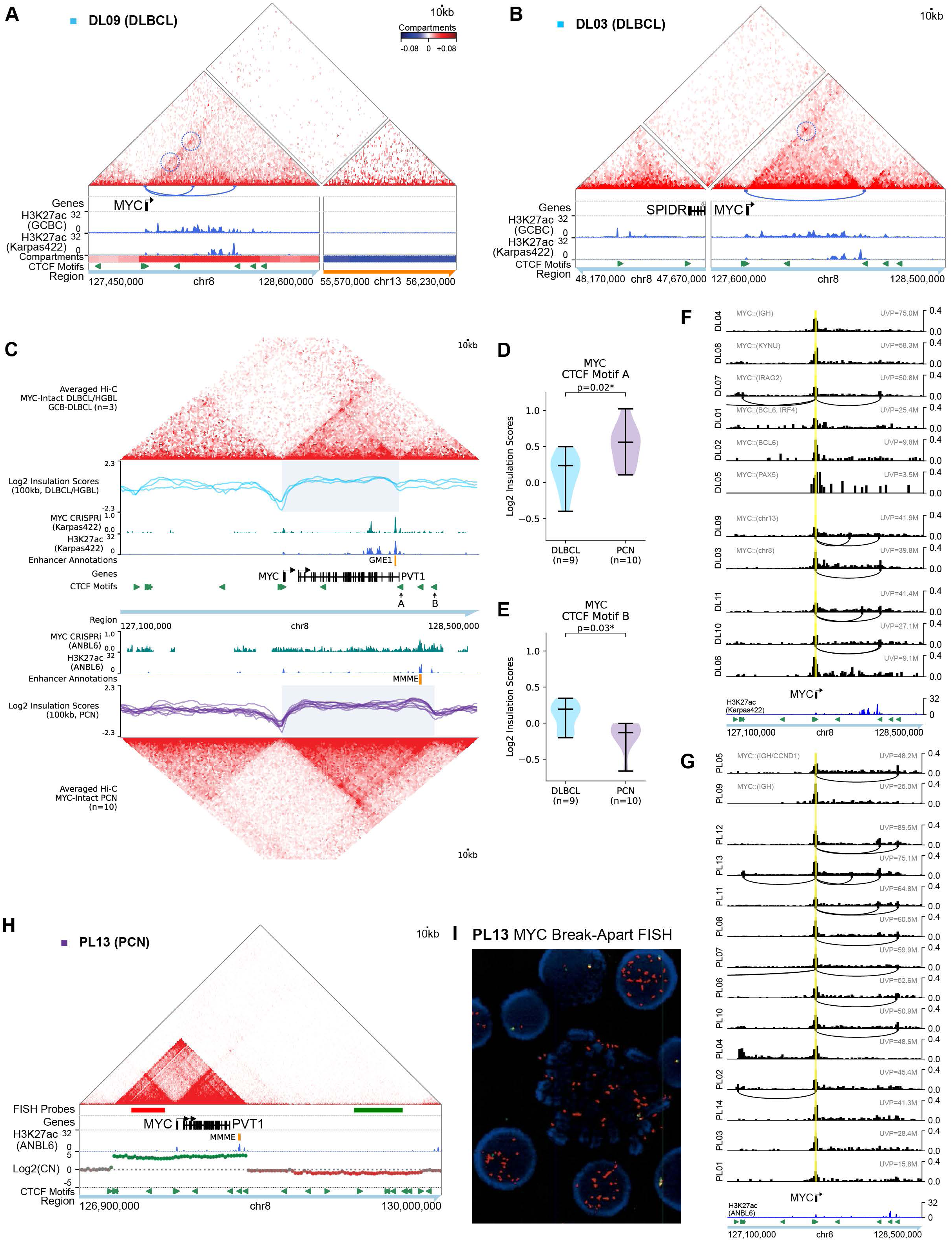
State-specific topology at the *MYC* locus. **A,** Balanced Hi-C matrix at 10kb resolution showing reconstruction of a rearrangement between *MYC* and a gene desert region of chr13 in DLBCL biopsy DL09. Significant loops involving the *MYC* promoter within its native locus are circled and shown as arcs. **B,** Balanced Hi-C matrix at 10kb resolution showing reconstruction of an intra-chromosomal fusion of *MYC* with the *SPIDR* locus in the context of chromosome 8 chromothripsis in DLBCL biopsy DL03. Loops involving the *MYC* promoter at its native locus are circled and shown as arcs. Note that reliable A/B compartment scores could not be calculated for chr8 in this dataset as chromothripsis-related heterologous interactions dominated eigenvector values. **C**, Top: Averaged balanced Hi-C matrices for *MYC*-intact GCB-DLBCL biopsies (n=3; DL06, DL10, DL11) at 10kb resolution across the *MYC* / *PVT1* locus. Aligned tracks include log2 insulation scores derived from the individual Hi-C samples calculated at 10kb resolution with 100kb windows, CRISPRi score for a high-throughput screen in the GCB-DLBCL cell line Karpas-422 (-log2 sgRNA depletion, 20-sgRNA sliding window, negative values not shown), and H3K27ac ChIP-Seq signal in Karpas-422. Bottom: Averaged balanced Hi-C matrix and log2 insulation scores as above, but for MYC-intact PCN without *MYC* amplification (n=10; PL01, PL02, PL03, PL06, PL07, PL08, PL10, PL11, PL12, PL14). CRISPRi score and H3K27ac ChIP-Seq signal are from the MM cell line ANBL6. Positions of the most essential enhancers identified by CRISPRi in Karpas-422 (GME-1) and ANBL6 (MMME) are indicated. Top and bottom: Gray boxes indicate 3’ *MYC* interacting regions demarcated by differential insulation boundaries at CTCF sites “A” and “B” in GCB-DLBCL versus PCN respectively. **D-E**, Violin plots showing the distribution of log2 insulation scores in *MYC*-intact systemic and CNS DLBCL (n=9; DC01, DC02, DC03, DC04, DC05, DC06, DL06, DL10, and DL11) and PCN (n=10, samples as in panel C) at the indicated CTCF motifs in panel C. P-values from statistical testing between the two groups are shown (Mann-Whitney U test). **F-G**, Virtual 4C analyses at 10kb resolution using the *MYC* promoter as the viewpoint (highlighted in yellow) across DLBCL samples (F) and PCN samples (G). Datasets are grouped by *MYC* rearrangement status (labelled by rearrangement partner at left) and ordered in descending order of unique valid pairs within each group (labelled at right). Loops detected from genome-wide Hi-C loop detection with HiCExplorer are shown as arcs. **H**, Raw Hi-C matrix at 10kb resolution showing a *MYC* amplification in PL13. Hi-C derived copy number is shown below. Genomic position of Vysis LSI *MYC* break-apart probes are shown in orange and green. Also shown are H3K27ac ChIP-Seq signal from MM cell line ANBL-6 and position of MM *MYC* enhancer (MMME) as in panel C. **I,** *MYC* break-apart FISH image for sample PL13, showing numerous separated red (5’) probes in each nucleus, consistent with extrachromosomal circular DNA.

### Topology supports role of state-specific *MYC* enhancers in DLBCL and MM

*MYC* rearrangements are associated with increased *MYC* expression in DLBCL and MM, but with significant overlap between rearranged and non-rearranged cases ^74,75^. We recently used CRISPR-interference in cell line models to functionally characterize a 3’ *MYC* locus enhancer that sustains *MYC* expression in non-*MYC* rearranged GCB-DLBCL (germinal center *MYC* enhancer 1; GME-1) ^71^ and a more distal 3’ enhancer that sustains *MYC* in non-*MYC* rearranged MM (multiple myeloma *MYC* enhancer; MMME ^76^). *MYC* activation by tissue-specific enhancers in epithelial cancers has been shown to be mediated in part by differential docking between a CTCF site at the *MYC* promoter and CTCF sites near tissue-specific enhancers ^77^. Consistent with this mechanism, FFPE Hi-C data showed an interaction domain (TAD) extending from the *MYC* promoter to an insulation boundary at a CTCF site (“A”) just distal to the GME-1 enhancer in non-MYC-rearranged GCB-DLBCL biopsies. In contrast, non-*MYC*-rearranged PCN datasets showed a different insulation boundary at a more distal CTCF site (“B”), which extended the *MYC* TAD to include the MMME (**Figure 6C-E** and **Supplementary Figure S6F-H**). Virtual 4C analysis also supported a different pattern of looping interactions between the *MYC* promoter and 3’ CTCF sites, with most *MYC*-intact GCB-DLBCL biopsies showing significant loops to CTCF site A / GME-1 (**Figure 6F**), while most *MYC*-intact PCN datasets showed significant loops involving the more distal site B (**Figure 6G**). Interestingly both of the DLBCL biopsies with *MYC* rearrangements that lacked expected features of enhancer hijacking (DL03 and DL09) showed strong looping of *MYC* to CTCF site A / GME-1, while this loop was only detectable in 1 of 6 biopsies with *MYC* rearrangement to proven or likely enhancer-donating loci. Hi-C also identified a case of MM (PL13) with a high-level *MYC* amplification that lacked evidence of juxtaposition to any external locus, but included the MMME in the amplicon (**Figure 6H**). *MYC* break-apart FISH in this case showed many separate copies of the centromeric (red) probe in each tumor cell (**Figure 6I**), suggesting that this lesion most likely represents extrachromosomal circular DNA containing the *MYC* gene and its native MMME enhancer, rather than an enhancer-hijacking rearrangement to an external locus.

## Discussion

In this study, we have shown that a genome-wide proximity method, Hi-C, can sensitively identify diagnostically and clinically significant genomic rearrangements in FFPE tissue across a range of lymphoid neoplasms, providing a potential alternative to low-throughput FISH assays. Lymphoid neoplasms may be particularly well-suited to this approach due to their high rate of clinically and diagnostically significant non-coding rearrangements, typically high tumor cellularity, and low rate of chromosomal instability compared to many epithelial cancers^78^.

The most obvious advantage of Hi-C over FISH is its genome-wide nature, which allows for detection of unanticipated oncogene rearrangements. Several of the unanticipated findings in our cohort were of significance for diagnostic classification, prognosis, and / or possible therapeutic prediction, and illustrate the advantage of a comprehensive approach over limited FISH panels. Identification of *BCL2* rearrangements is crucial for diagnosis of “High-grade B-cell lymphoma with *MYC* and *BCL2* rearrangements” (HGBCL–DH–BCL2), a poor-prognosis variant of DLBCL that is often treated with more intensive chemotherapy regimens^15,16^ than the R-CHOP regimen typically used for DLBCL. Many clinical labs detect these lesions with *IGH*::*BCL2* dual fusion FISH probes, since this rearrangement is specific for follicular lymphoma and GCB-DLBCL, while other *BCL2* alterations such as gene amplification do not carry the same diagnostic associations^18,61^. However this strategy can fail to detect rare rearrangements between *BCL2* and alternate activating partners, such as the *BCL2*::*IGL* rearrangement we identified by Hi-C in DL03, which could result in failure to identify HGBCL–DH–BCL2 and subsequent undertreatment.

Testing for *CCND1* rearrangement (by FISH or high-specificity immunohistochemistry) is routinely performed in mantle cell lymphoma, where it plays a key role in diagnostic classification^1,2^, and in MM, where it can contribute to therapeutic risk stratification^7^. Rearrangement of the homologous *CCND2* gene, an unanticipated finding of Hi-C in our cohort for case PL08, is not routinely tested in most centers due to its rarity, despite this lesion having similar biological function. A similar situation applies to rearrangements of the *MAF* gene and its homologs, which have important implications for prognosis and therapeutic stratification in MM^6,7^. While the expanded MM FISH algorithm now in place at NYU would likely have detected the unanticipated *IGH::MAFB* rearrangement found by Hi-C in case PL07, this is not true for other variants, such as *MAFA* rearrangements and *MAF* gene rearrangements to non-*IGH* partners.

The two rearrangements involving PD1 ligand gene loci further demonstrate the potential clinical value of using an unbiased genome-wide approach for rearrangement detection. Checkpoint inhibitor therapy has proven benefits in classic Hodgkin lymphoma ^79^ and primary mediastinal large B-cell lymphoma ^57,58^, which activate PD-ligand genes by multiple mechanisms, including JAK signaling and frequent SVs of the PD ligand gene locus on 9p24.1 such as long-distance rearrangements and tandem amplifications. The PD-L2 gene rearrangement we identified in case PM01 therefore supports the diagnosis of PMBL, for which a checkpoint inhibitor such as pembrolizumab or nivolumab is recommended in second-line therapy ^80^. Structural abnormalities of the PD ligand gene locus are also common in primary CNS DLBCL ^81^, a rare disease with very poor prognosis when refractory to first-line therapy. Clinical responses to pembrolizumab and nivolumab have been reported in PCNSL ^82–84^, and several clinical trials of these agents are ongoing^85,86^. Our identification of an apparent PD-L1-activating rearrangement by Hi-C in PCNSL biopsy DC05 could therefore motivate participation in such a trial. PD ligand gene rearrangements are rarer in systemic DLBCL, but do occur ^18,68^, and our findings suggest that Hi-C could be an effective method to identify such patients.

Another key advantage of Hi-C is the structural detail it provides with regard to genomic breakpoint position and the identity of oncogene rearrangement partner loci. Unlike sequencing of coding mutations, FISH does not precisely define the nature of the genetic lesion, and interpretation relies on established knowledge about the diagnostic or clinical associations of common rearrangements at the targeted loci. However, these assumptions can be problematic for loci that can undergo multiple types of rearrangements with different functions, or that show increased rates of functionally irrelevant DNA damage, both of which are true for key rearrangement loci in lymphoid cancers such as the immunoglobulin and *BCL6* loci ^87,88^. Although Hi-C also does not provide nucleotide-level resolution, identifying fusion position on the order of kilobase to 10-kilobase resolution, these details were sufficient in our cohort to distinguish between promoter swap rearrangements that activate *BCL6*, rearrangements that “donate” the *BCL6* LCR enhancer to activate *MYC*, and other rearrangements (e.g. those seen in DL10 and DL03) that did not fit either functionally characterized pattern. Although break-apart FISH is commonly used to identify *MYC* rearrangements, there is substantial evidence to indicate that all *MYC* rearrangements are not equivalent in their clinical and biological implications. *IG*::*MYC* rearrangements uniformly result in high *MYC* expression and confer a poorer prognosis in DLBCL, while non-*IG*::*MYC* rearrangements are heterogeneous in their effects on MYC expression and as a group do not show the same prognostic effect as *IG*::*MYC* rearrangements ^13,89^. Similarly, *MYC*::*IGL* rearrangements appear to be associated with a worse prognosis in MM than MYC rearrangements to other partners, further highlighting the importance of partner identification ^90,91^.

A final advantage of Hi-C is the ability of genome topology details such as A/B compartment state, insulation boundaries, loops between CTCF sites, and enhancer-promoter interactions to inform the interpretation of rearrangement function, which we demonstrated for common recurrent rearrangements, and propose as helpful clues for interpreting function for unfamiliar combinations of loci. For example, we are only aware of a single prior report of rearrangement between *MYC* and the *KYNU* locus^30^, and no prior reports for *MYCN* and the *ARHGAP24* locus, but the presence of genome-wide significant neo-loops in these datasets between the oncogene and candidate partner locus enhancers (based on reference H3K27ac ChIP-Seq data) helped support these as likely functional events. Biopsies DL09 and DL03 provided examples of the reverse situation, where topological interaction details provided by Hi-C reduced confidence that *MYC* rearrangements (originally identified by break-apart FISH) were functional oncogene-activating lesions. In these biopsies, statistically significant physiological loops were present between the *MYC* genes and native 3’ distal enhancer regions (supporting the quality of topology data in these biopsies), but there was no evidence of heterologous neo-loops to between the *MYC* gene and rearrangement partner loci, with the DL09 partner locus showing a uniform B compartment (heterochromatin) state.

While our study highlights the promise of Hi-C for detection of oncogenic rearrangements in clinical samples, some important limitations are also evident. Unlike WGS, which detects SVs via direct mapping of read-pairs across genomic fusions, sensitivity of Hi-C for SV detection can be directly affected by the size of altered region. Thus, while automated rearrangement detection algorithms like Hi-C Breakfinder and EagleC are sensitive for intergenic rearrangements and large intragenic SVs such as the recurrent ∼3 Mb deletion that fuses the *RAG2* and *LMO2* loci, smaller deletions such as the recurrent *STIL*::*TAL1* fusion may not be resolved with current algorithms. Further work is needed to determine the sensitivity of Hi-C for small genomic insertion events on the order of ∼100 kb, which are known to occasionally result in enhancer-hijacking activation of oncogenes such as *MYC* and *BCL2* while remaining cryptic to FISH ^92^. Detection of such events by Hi-C would likely be more sensitive to effective sequencing depth than fusion of large, contiguous genomic regions. Our cohort also highlighted limitations for detection of subclonal events, such as the *TRA::MYC* rearrangement in case BA01, and this limitation would likely also apply to specimens with a high fraction of non-neoplastic tissue.

An important requirement for implementation of Hi-C in routine clinical diagnostics will be standardization of analysis and reporting approaches. The resolution provided by Hi-C is superior to FISH, but the lack of nucleotide-level resolution of breakpoints may result in ambiguity of exons included in some gene fusions. More challenging will be reporting of non-coding rearrangements. One approach, analogous to the tier system used for molecular reporting of coding mutations ^93^ would be to classify potential enhancer-hijacking events near known recipient oncogenes into tiers based on prior experience, with rearrangements involving functionally characterized or highly recurrent enhancer-donating loci classified as high-confidence events, while rearrangements involving previously unreported partner loci could be classified as events of uncertain significance. Recent efforts to comprehensively map recurrent rearrangement partners of common DLBCL oncogenes, for example, will provide a valuable resource ^29^. Our findings indicate that topology information may assist in the interpretation of candidate enhancer-hijacking events due to the present or absence of neo-loops, active compartment state of the partner locus, or presence of candidate active enhancers in appropriate reference chromatin datasets. Further work is needed to standardize and validate the utility of identifying such features for routine clinical diagnostics.

## Materials and Methods

### Study Cohort

We profiled 44 lymphoid cancer biopsies retrospectively from pathology departments across three institutions (NYU Langone Health, n=22; Michigan Medicine, n=20; Massachusetts General Hospital, n=2), including archival FFPE biopsies up to 17 years old. Cohort demographics, formal clinical diagnoses and known genomic findings from prior clinical testing with conventional cytogenetics, DNA microarray, and FISH were obtained from clinical records and are detailed in **Supplementary Table S1** and **Supplementary Table S2.** This study was performed in accordance with the Institutional Review Boards (IRB) of NYU Langone (IRB#: i14-00948) and the University of Michigan Medical School (HUM00155777).

Other datasets used included Hi-C from hematological (GM12878, KBM7, K562) and non-hematological (HMEC, HUVEC, IMR90, NHEK) cell lines from GEO Series GSE63525 ^38^ and for patient-derived plasma cells (PC), memory B cells (MBC), germinal center B cells (GCB) and naïve B cells (NBC) from the EGA dataset EGAD00001006485 ^39^. Oriented CTCF motifs were previously defined ^105^. H3K27ac datasets for plasma cells, germinal center B-cells, naïve B cells, and a primary T-ALL sample were obtained from the BLUEPRINT project ^98^. H3K27ac and MYC CRISPRi for a DLBCL cell line (Karpas422) and a MM cell line (ANBL6), as well as GRCh38 coordinates of tissue-specific enhancer annotations, were previously published ^68,76^.

### Sample Preparation and Sequencing

Samples underwent Hi-C library preparation and sequencing at Arima Genomics. FFPE tissue from 2-5 unstained slides cut at 5-10µm were processed using the Arima HiC+ for FFPE kit (Product Number A311038, Arima Genomics, Carlsbad, CA) as per manufacturer protocols to produce Illumina-compatible sequencing libraries. Briefly, 5µm unstained FFPE tissue sections were first de-waxed and rehydrated, and then subjected to Hi-C sample preparation using Arima HiC+ for FFPE kit. Following Hi-C sample preparation, Illumina-compatible sequencing libraries were constructed by shearing the proximally ligated DNA and then size selecting DNA fragments using SPRI beads. The size-selected DNA fragments containing proximity ligation junctions were then enriched using Enrichment Beads (provided in the Arima HiC+ for FFPE kit) and converted to Illumina-compatible sequencing libraries. The resulting Hi-C libraries underwent standard quality control (qPCR and BioAnalyzer) and were sequenced on Illumina NovaSeq 6000 as per manufacturer protocols to a depth between 101 million and 536 million raw read pairs.

### Generation of Hi-C Matrices

Hi-C sequencing data was processed using the Arima-SV-Pipeline (v1.3; https://github.com/ArimaGenomics/Arima-SV-Pipeline), comprising HiCUP (v0.8.0) ^94^ to calculate quality control metrics and perform read alignment and filtering, hic_breakfinder ^35^ to call rearrangement breakpoints, and Juicer Tools (v1.6) ^95^ to produce multi-resolution Hi-C matrices from mapped and filtered read pairs. FFPE samples were processed using default pipeline parameters for the Arima-HiC+ for FFPE restriction enzyme chemistry; reference non-FFPE samples were processed as described above, except using modified digest and cut site files for the respective enzyme (MboI or DpnII) that were produced by HiCUP Digester and the generate_site_positions utility available from Juicer respectively. Alignment was performed against the GRCh38 human reference genome.

### Identification of Topological Features

Chromosomal A/B compartments were identified by calculating compartment scores separately for each autosomal chromosome at 100kb resolution using the eigenvector utility from Juicer Tools (v1.6). To ensure the signs of compartment scores followed a convention of positive signs corresponding with active chromatin states ^96,97^, the signs of compartment scores for an entire chromosome were flipped if the Pearson correlation with reference H3K27ac enrichment data from corresponding cell types ^98^ was initially negative (after masking compartment regions overlapping with segmental duplications ^99^ and ENCODE blacklist regions in GRCh38 ^100^).

Insulation scores were calculated using cooltools (v0.7.1) ^101^ at 10kb resolution using 100kb windows, and the gradient of insulation scores was calculated using the gradient function from numpy (v1.26.4). Hierarchical TADs were called from Hi-C data using TADlib (v0.4.5-r1) ^102^ at 25kb resolution. Loops were called using the hicDetectLoops utility from HiCExplorer (v3.7.3) ^103^ at 10kb resolution with a maximum loop distance of 5Mb.

### Detection of Structural Variants and Neo-loops

To increase our sensitivity for detecting genomic rearrangements, we combined the results of two different structural variant callers to produce a merged set of candidate rearrangement breakpoints for each sample. In addition to breakpoint calls from hic_breakfinder, we used EagleC (v0.1.9) ^104^ on Hi-C matrices converted to multi-resolution cool format using HiCExplorer (v3.7.3) ^103^ to independently identify rearrangement breakpoints and subsequently merged hic_breakfinder and EagleC breakpoint calls by union. Where hic_breakfinder and EagleC made calls with both anchors within 1Mbp proximity of each other, we selected the EagleC breakpoint for the merged set. We manually reviewed the Hi-C matrix data and all breakpoint calls in this merged call set to ensure breakpoints were located at precise Hi-C signal boundaries with evidence of proximity-based signal decay in the direction of the breakpoint strands. Further details of manual review are included in the **Supplementary Methods**.

Copy number was calculated from Hi-C data at 25kb resolution using the calculate-cnv and segment-cnv utilities from NeoLoopFinder (v0.4.3-r2) with default parameters ^37^. Copy number and structural variant calls were used to normalize and reconstruct local chromosomal assemblies around rearrangement breakpoints, and *de novo* chromatin loops forming across rearrangement breakpoints (“neo-loops”) were called from Hi-C data using NeoLoopFinder (v0.4.3-r2) ^37^ at a probability threshold of 0.90.

### Feature Enrichment

Enrichment of TAD boundaries and loop anchors for CTCF motifs was assessed by counting the number of features (within 1 bin (±25kb) of called TAD boundaries or within the called loop anchor width respectively) containing CTCF motifs from public reference data ^105^. Aggregate peak analysis (APA) was performed at loop and neo-loop anchors by aggregating Hi-C signal (balanced using square root vanilla coverage and normalized by unique valid pairs) at 10kb resolution within 100kb of all loop/neo-loop anchors for anchors with a minimum distance of 300kb. Neo-loop anchor enrichment for active enhancers was assessed by calculating mean H3K27ac ChIPSeq binding signal (from public reference germinal center B-cells) occurring within 100kb of loop/neo-loop anchors across the sample cohort for loop/neo-loop anchors at least 100kb from chromosomal endpoints. Enrichment of TAD boundaries and loop/neo-loop anchors were compared against an equal number of random genomic loci matched for the same chromosomes as TAD boundary and loop/neo-loop calls. Compartment pair enrichment for neo-loops was calculated by determining the frequency of neo-loop anchors occurring in compartments with positive (for active) and negative (for inactive) compartment scores, and compared against random genomic loci drawn from the same chromosomal distribution as neo-loop anchors.

### Visualization of Hi-C Data

Hi-C data was visualized with custom plotting code in matplotlib in conjunction with hic-straw (v1.3.1) to access Hi-C data values in Python. Unless otherwise specified, Hi-C matrices are visualized using the observed values balanced by weight vectors from Juicer’s “fast scaling” algorithm (where balancing weights successfully converged) or square root vanilla coverage. Gene tracks show Ensembl v110 gene annotations for GRCh38 accessed via pyensembl (v2.3.13). Virtual 4C tracks were generated from Hi-C data by extracting a subset of Hi-C values at a fixed viewpoint on one axis (analogous to taking a “column” of Hi-C matrix data); where specified to address matrix sparsity at high resolution, a sliding window of ±1 bin (at the virtual 4C resolution) was taken around the fixed viewpoint and Hi-C matrix values across the fixed viewpoint window were averaged. Virtual 4C comparisons of observed Hi-C values across different samples are normalized by the unique valid pairs in each sample. Circos plots were created using pyCirclize (v.1.6.0).

### Comparative and Statistical Analyses

Principal component analysis of compartments scores (excluding compartments overlapping with blacklisted GRCh38 regions) was performed using scikit-learn (v1.4.2). Hi-C matrix data and topological feature calls for each sample were compared for similarity against a high-quality reference Hi-C dataset from the GM12878 cell line. The similarity of TADs and loops between individual samples was calculated by binning TAD boundaries and loop anchors into 50kb bins respectively and subsequently calculating the Jaccard similarity statistic between the sets of TAD boundaries and loop anchors between each pair of samples. Comparisons of compartment and insulation scores at a specific locus were conducted using Mann-Whitney U tests, while comparisons of TAD boundaries and loop anchors at specific loci were conducted using Fisher’s exact test on contingency tables representing the presence or absence of a TAD boundary or loop anchor in 100kb and 50kb genomic bins respectively amongst samples with more than 10 million unique valid pairs. In both instances, significant feature differences were defined at a p-value threshold of <0.05.

### Data Availability

The processed Hi-C matrices generated from samples in this study have been deposited to the Gene Expression Omnibus (GEO) (GSEXXXXXX, and four samples previously deposited in GSE230396).

## Supporting information

Supplementary Test and Figures

## Authors’ Contributions

**J. Wu**: Data curation, formal analysis, methodology, software, visualization, writing – original draft, writing – review & editing. **A. Chu**: Data curation, formal analysis, methodology, software, visualization, writing – original draft, writing – review & editing. **J. Cho**: Data curation, investigation, validation, writing – review & editing. **M. Movahed-Ezazi**: Data curation, formal analysis, investigation, writing – review & editing. **K. Galbraith**: Data curation, investigation, writing – review & editing. **C. Fang**: Data curation, investigation, writing – review & editing. **Y. Yang**: Data curation, software, writing – review & editing. **C. Schroff**: Data curation, writing – review & editing**. K. Sikkink**: Data curation, investigation, writing – review & editing. **M. Perez-Arreola**: Investigation, writing – review & editing. **L. Van Meter**: Investigation, writing – review & editing. **S. Gemus**: Investigation, writing – review & editing. **J. Belton**: Data curation, writing – review & editing. **V. Nardi**: Data curation, resources, writing – review & editing. **A. Louissant Jr**: Data curation, resources, writing – review & editing. **D. Shasha**: Conceptualization, formal analysis, software, supervision, writing – review & editing. **A. Tsirigos:** Data curation, formal analysis, resources, supervision, validation, writing – review & editing. **N. Brown**: Investigation, validation. **T. Gindin**: Conceptualization, data curation, formal analysis, resources, supervision, validation, writing – review & editing. **M. P. Cieslik**: Conceptualization, formal analysis, supervision, writing – review & editing. **M. Kim**: Conceptualization, formal analysis, methodology, supervision, validation, writing – review & editing. **A. Schmitt**: Conceptualization, data curation, formal analysis investigation, methodology, resources, writing – original draft, writing – review & editing. **M. Snuderl**: Conceptualization, data curation, formal analysis, funding acquisition, investigation, methodology, project administration, resources, supervision, validation, visualization, writing – original draft, writing – review & editing. **R. J. H. Ryan**: Conceptualization, data curation, formal analysis, funding acquisition, investigation, methodology, project administration, resources, supervision, validation, visualization, writing – original draft, writing – review & editing.

## Acknowledgements

This work was supported by NIH R01CA245059 (R.J.H.R), and in part by a grant from Ian’s Friends Foundation (M. S.), and NIH 5R01GM121753 and NYU Wireless (D. S.). A.T. is supported by the NCI/NIH P01CA229086, NCI/NIH R01CA252239, NCI/NIH R01CA260028 and NIH/NCI R01CA140729. M.K. is supported by the NIH R00HG011542. M.S. and C.S are supported by the National Institute of Neurological Disorders and Stroke grant R01-NS122987. We thank the authors Vilarrasa et al and V. Chapaprieta for liaison and provision of reference Hi-C data from normal blood cells. We thank Nina Murrell for coordinating data transfer of raw reads to NYU. The authors thank the NYU Langone High-Performance Computing Core for access and support for computational resources used during this study. The author (J.W.) gratefully acknowledges financial support from the Australian-American Fulbright Commission and The Kinghorn Foundation.

