## Supplementary Test and Figures for "Hi-C for genome-wide detection of enhancer-hijacking rearrangements in routine lymphoid cancer biopsies"

#### Supplementary Table S1

Table showing overview of pre-analytical cohort characteristics, including patient demographics, and biopsy age and sites.

#### Supplementary Table S2

Table showing formal diagnoses, known SVs, and corresponding selected SVs from Hi-C for each case. Cases reported in a prior publication [71] with an alternate sample ID are shown in brackets.

#### Supplementary Table S3

Table of quality control metrics for all samples (including reference samples). Quality metrics are obtained from the QC metric summary from the Arima-SV-Pipeline.

#### Supplementary Table S4

Table showing similarity metrics of each sample against GM12878 (Pearson correlation of compartment and insulation scores, Jaccard similarity of 50kb-binned TAD boundaries and loop anchors), corresponding with Fig 1A.

#### Supplementary Methods

##### Review of Merged Breakpoint Calls

All breakpoint calls were manually reviewed for the following criteria:

1. Breakpoints must show sharp increase in signal at both breakpoint anchors, resulting in a distinctive, right-angled "corner"-like appearance.
2. Breakpoint anchors must be situated precisely at the point of increase of signal.
3. Breakpoints must show evidence of proximity-based decay, i.e. Hi-C interactions should decrease with further distance away from the breakpoint anchors.
4. Breakpoint strands must match the direction of signal decay away from the breakpoint.
5. Breakpoint calls must not be present in the non-rearranged public reference data sets (as these were deemed more likely to be artifacts).

Breakpoints close to the main diagonal with anchors within 5Mb apart and consisting of +- strands were automatically excluded as these were deemed indistinguishable from TADs (except where coverage was exceptionally greater at this region, which was considered suggestive of amplification). Breakpoints which did not show sharp "corner"-like signal with evidence of distance decay were excluded. Breakpoints which were not positioned precisely at the "corner" of signals were adjusted to precisely match the boundary of signal increase. Breakpoints for which strandness calls did not match the direction of signal decay were adjusted for the correct strandness. Hi-C breakpoints which fulfilled the above criteria upon careful manual review, but were not called by the automated callers, were added to the final breakpoint set and are noted where applicable in the main body of the text.

#### Supplementary Figure S1

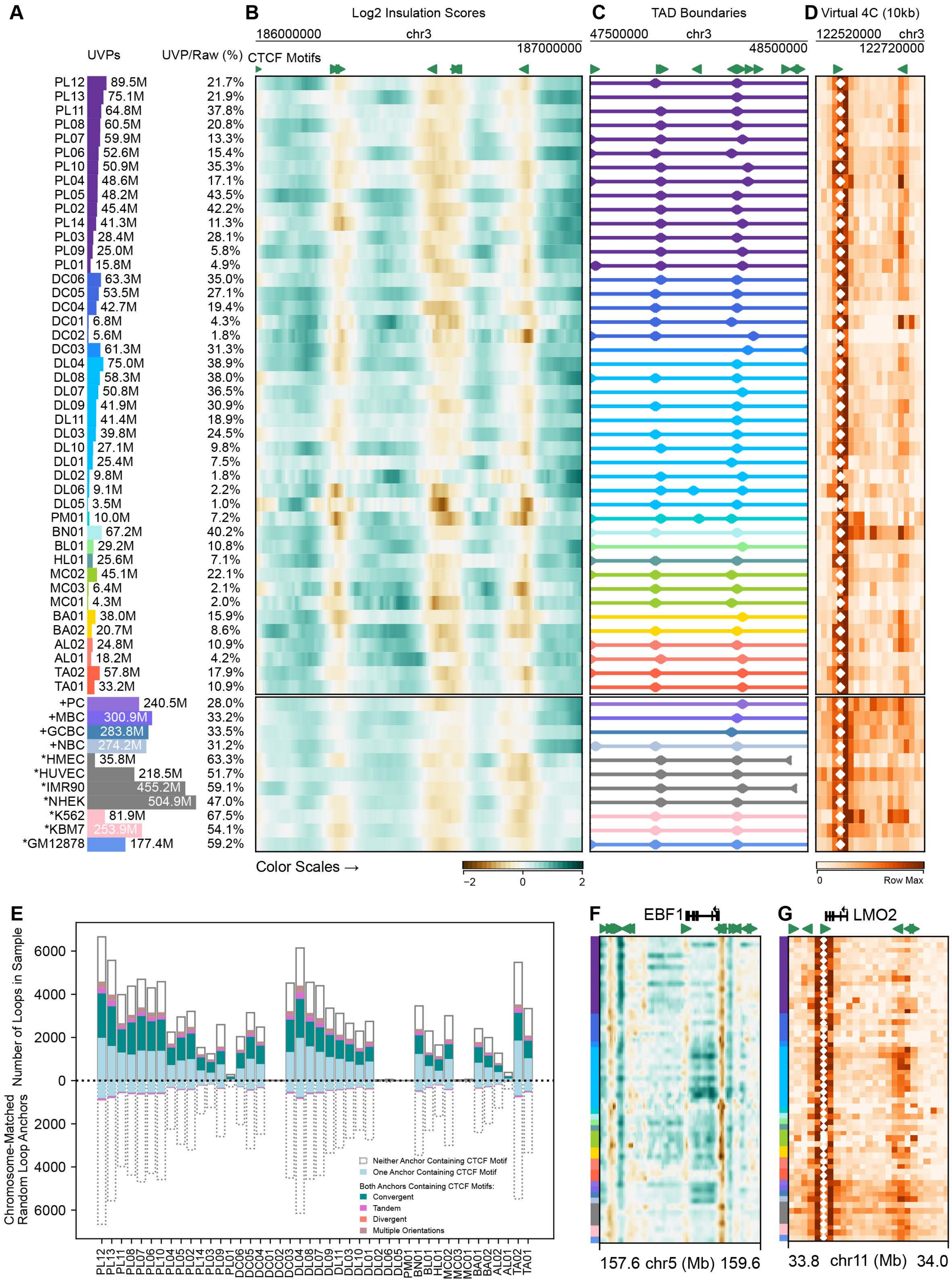

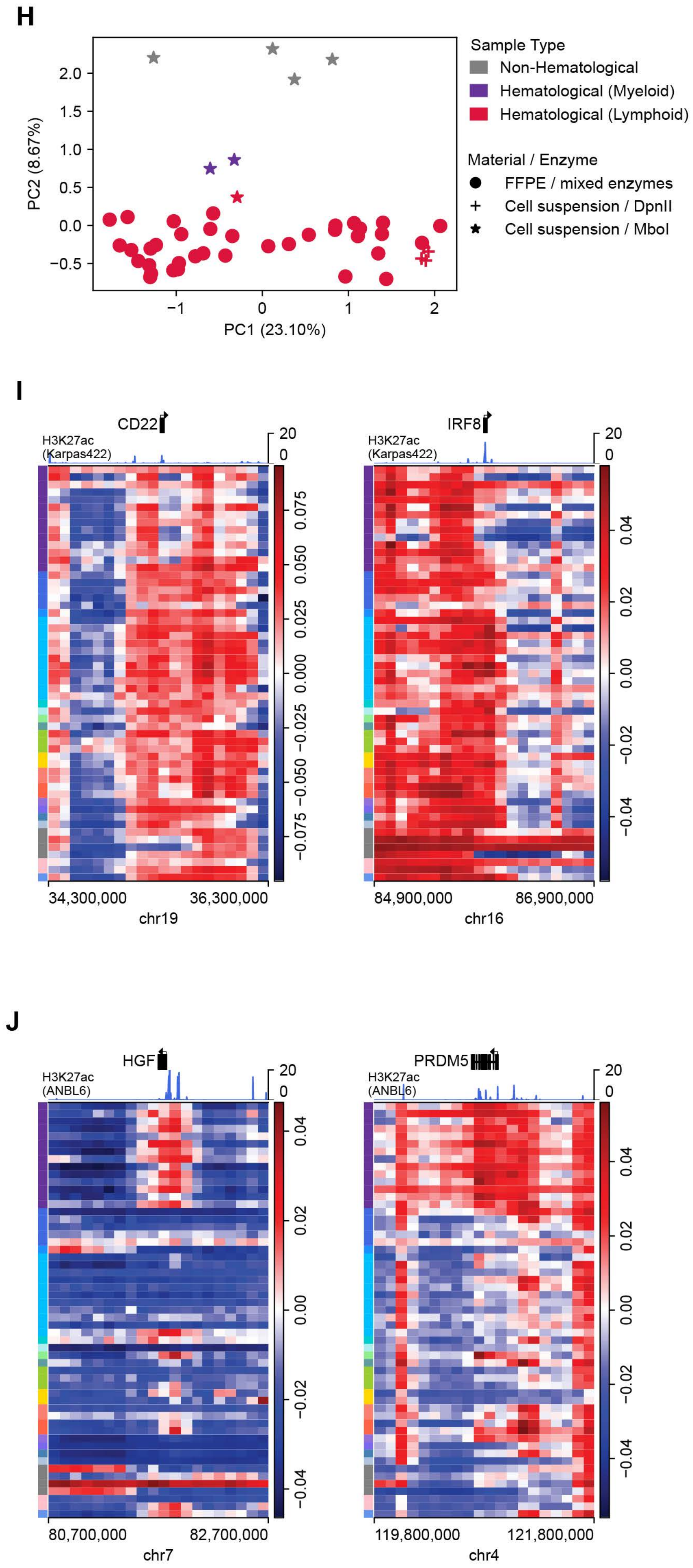
H
Sample Type
Non-Hematological
Hematological (Myeloid)
Hematological (Lymphoid)
Material / Enzyme
FFPE / mixed enzymes
Cell suspension / DpnII
Cell suspension / MboI
PC2 (8.67%)
PC1 (23.10%)
I
CD22
IRF8
H3K27ac
(Karpas422)
34,300,000
36,300,000
chr19
84,900,000
86,900,000
chr16
J
HGF
PRDM5
H3K27ac
(ANBL6)
80,700,000
82,700,000
chr7
119,800,000
121,800,000
chr4

**K**

Hi-C and TAD Calls for window chr5:157625000:159625000 (25kb)

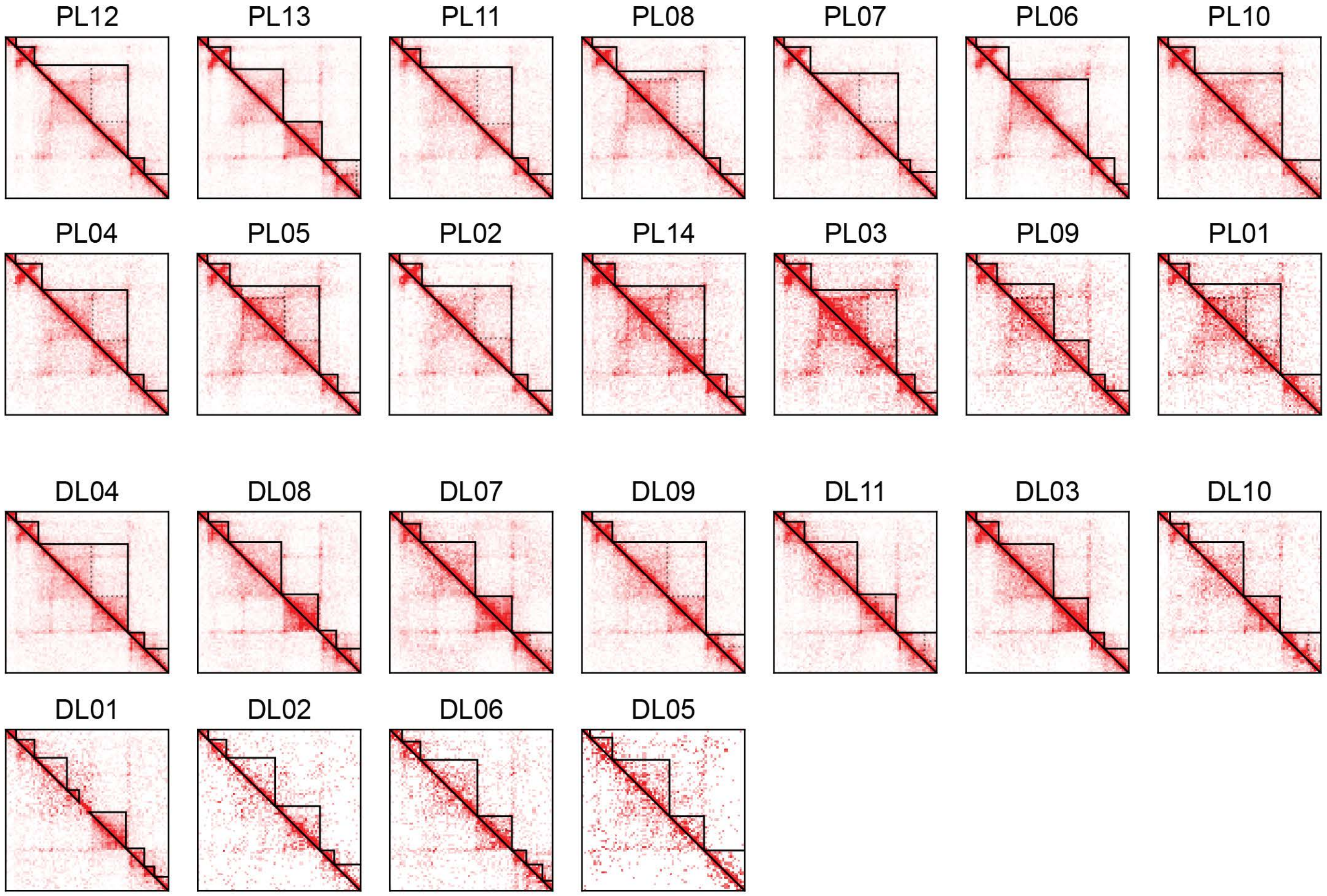


**L**

Hi-C and Loop Calls for window chr11:33710000:34110000 (10kb)

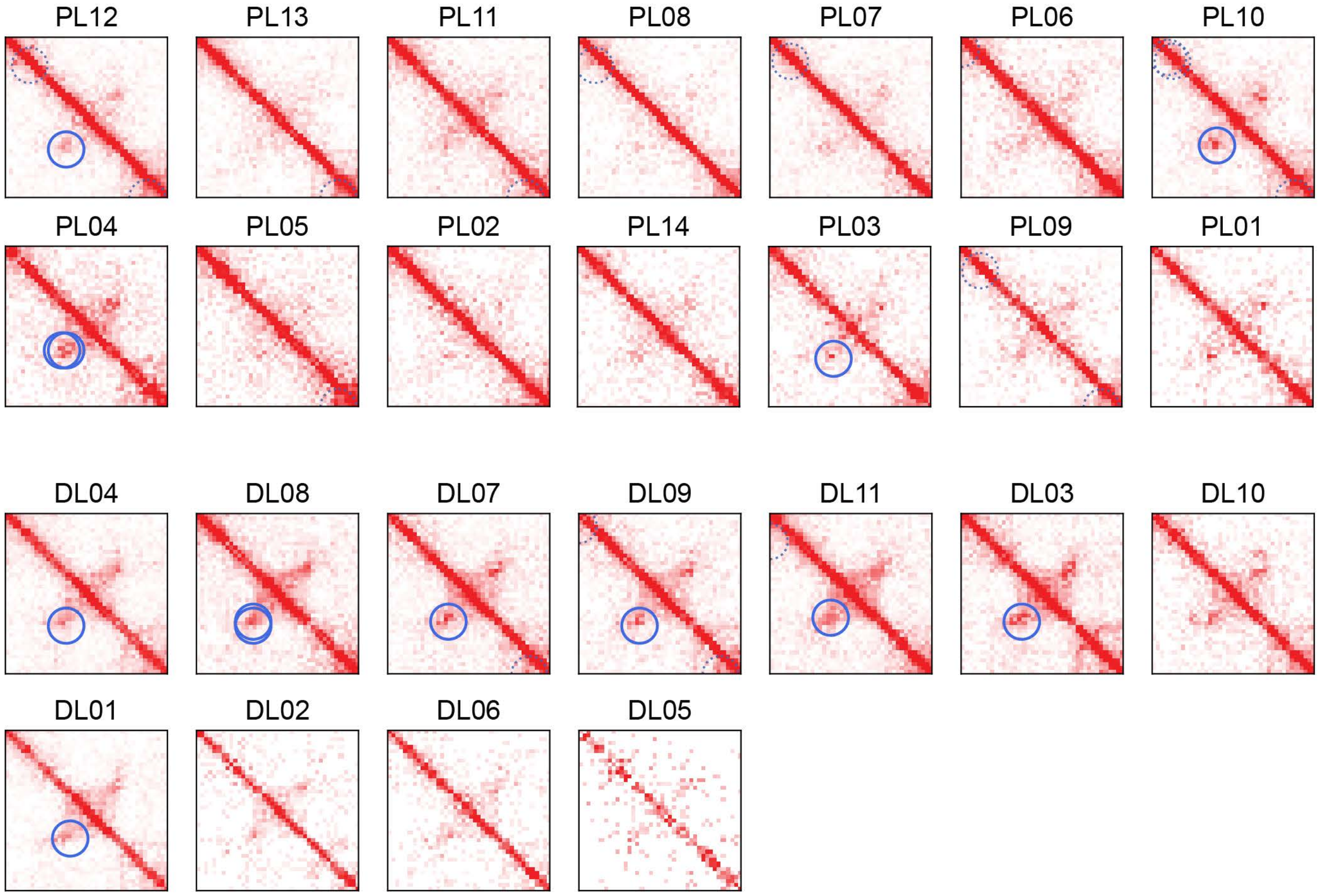

#### Supplementary Figure S1

**A,** Number of unique valid read-pairs (UVP) and yield (UVP/ raw read-pairs) for all samples in the FFPE and reference cohorts. Samples are color coded by disease type as in Figure 1

**B,** Representative region of chr3 showing Log2 insulation scores across the cohort. Note correlation of insulation boundaries with CTCF motifs at top.

**C.** Representative region of chr3 showing topologically-associating domain (TAD) boundaries across the cohort. Note correlation with CTCF motifs at top.

**D.** Representative region of chr3 showing virtual 4C interactions with a viewpoint (white diamonds) aligned to a CTCF motif at top

**E**, (Top) Stacked bar chart showing the presence and orientation of CTCF motifs (in 10kb resolution loop anchors) for all loops detected in each sample. (Bottom) CTCF motif statistics for an equal number of chromosome-matched random anchors for each sample.

**F**, Log2 insulation scores across the *EBF1* locus (corresponding with the region in Figure 1I). Samples are ordered and colored as in panel A.

**G**, Virtual 4C at 10kb resolution across the *LMO2* locus (corresponding with the region in Figure 1J). Viewpoint is indicated by white diamonds. Samples are ordered and colored as in panel A.

**H**, Plot of first two principal components over sample compartments across all samples. Samples are colored by broad category (non-hematological, myeloid and lymphoid) with symbols indicating material type and enzyme used.

**I**, Loci of developmentally regulated genes CD22 and IRF8 (downregulated in plasma cell compared to mature B cells) showing differential compartment states between mature B cell lymphomas and plasma cell neoplasms. Sample color mapping and ordering as in Figure 1A and Supplementary Figure S1A.

**J,** Loci of developmentally regulated genes HGF and PRDM5 (upregulated in plasma cell compared to mature B cells) showing differential compartment states between mature B cell lymphomas and plasma cell neoplasms. Sample color mapping and ordering as in Figure 1A and Supplementary Figure S1A.

**K,** Balanced Hi-C contact matrices at 25kb resolution showing hierarchical TAD calls for PCN and DLBCL in the region chr5:157,625,000-159,625,00 (corresponding with Figure 1I). The outermost TAD boundaries are shown in solid black; inner TAD hierarchies are shown as dotted lines.

**L**, Balanced Hi-C contact matrices at 10kb resolution showing loop calls for PCN and DLBCL samples in the region chr11:33,710,000-34,110,000 (corresponding with Figure 1J). Loops are circled in solid blue.

**Supplementary Figure S2**

**A**

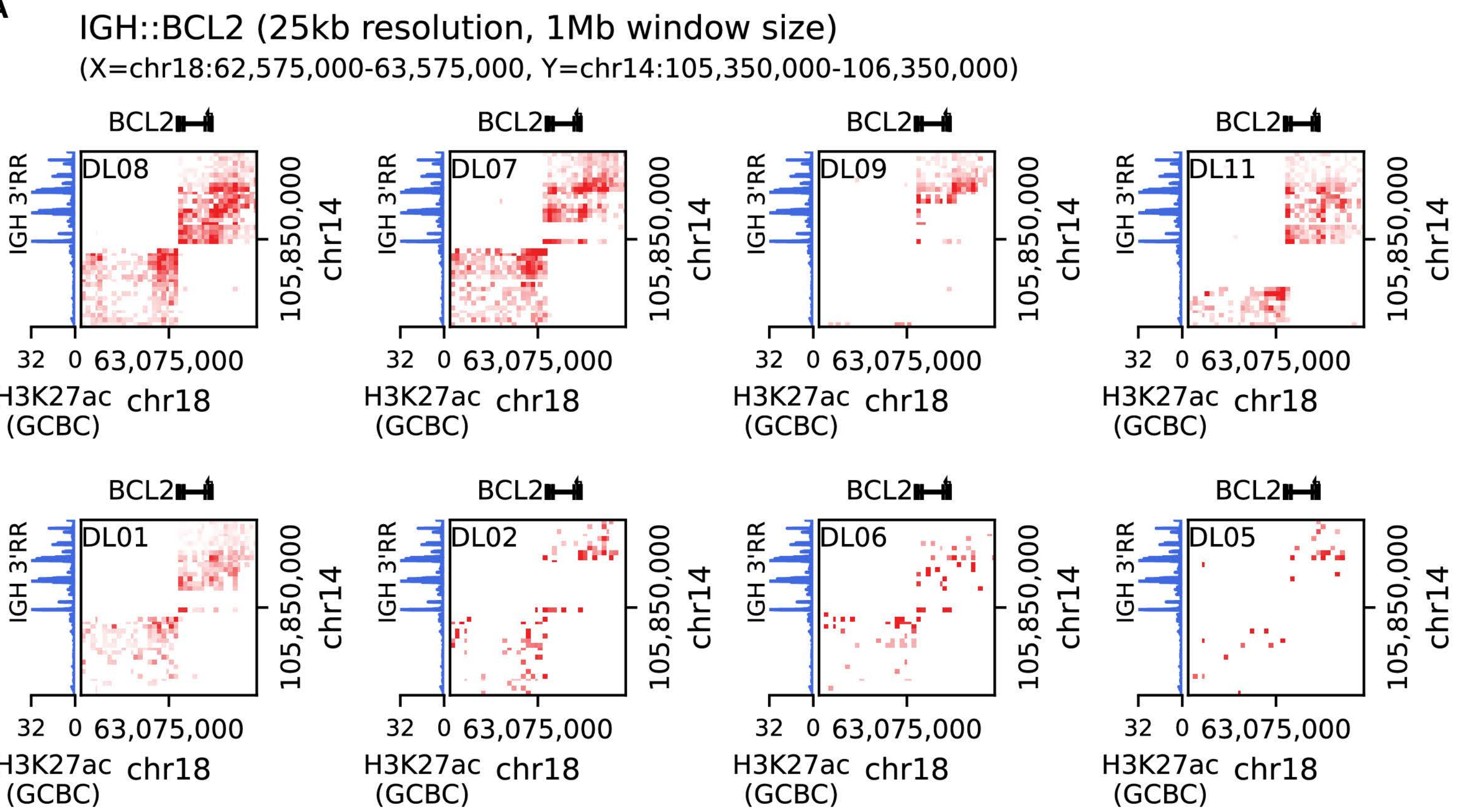


**B**

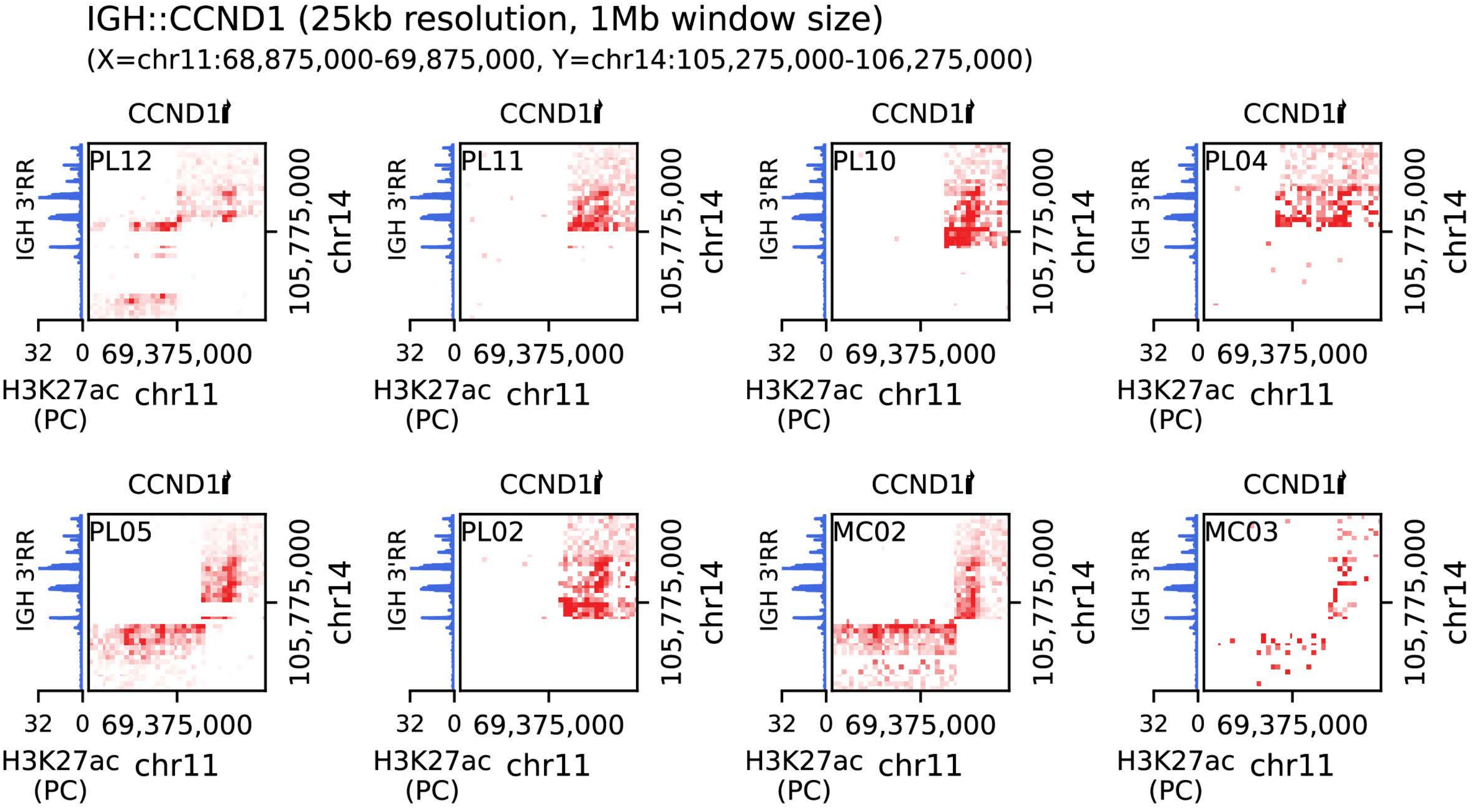

C

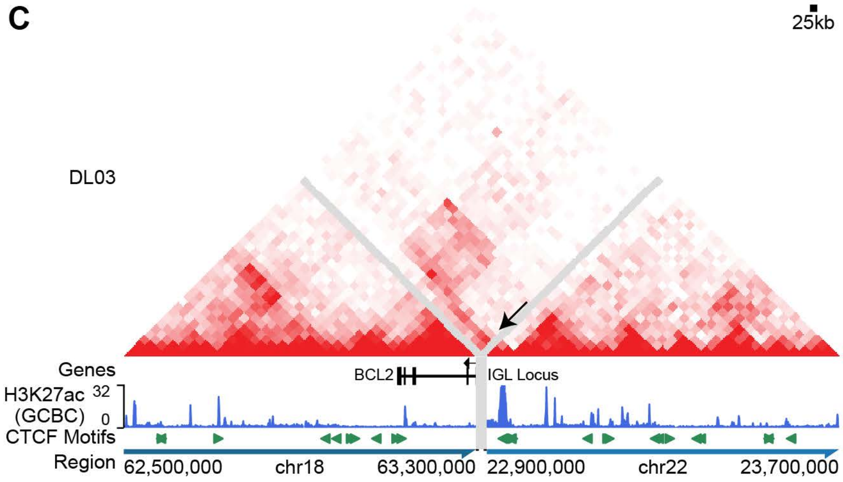
25kb
DL03
Genes
BCL2
IGL Locus
H3K27ac
(GCBC)
32
0
CTCF Motifs
Region
62,500,000
chr18
63,300,000
22,900,000
chr22
23,700,000

D

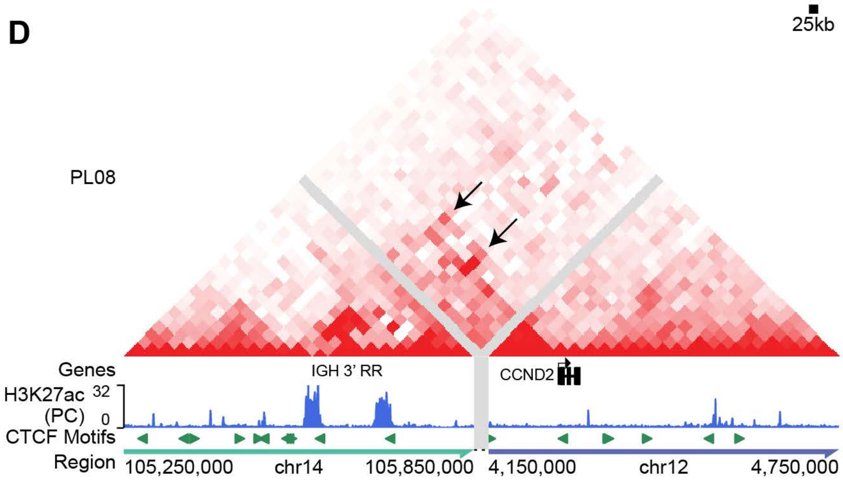
25kb
PL08
Genes
IGH 3' RR
CCND2
H3K27ac
(PC)
32
0
CTCF Motifs
Region
105,250,000
chr14
105,850,000
4,150,000
chr12
4,750,000

E

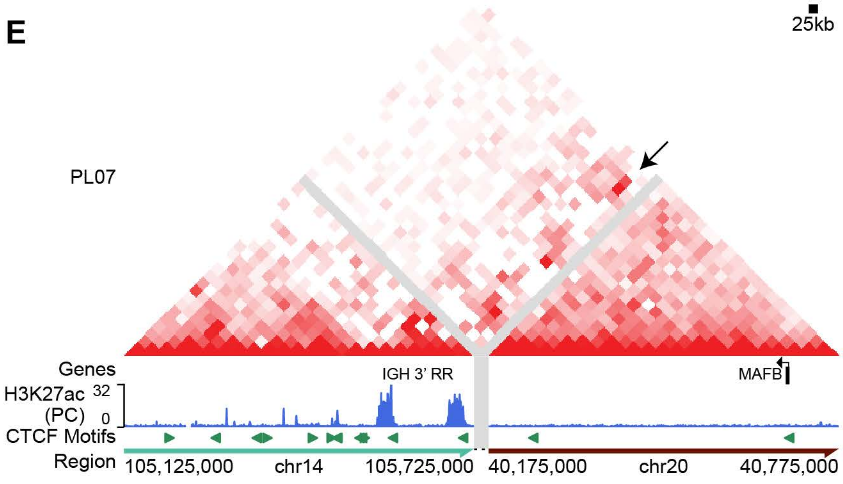
25kb
PL07
Genes
IGH 3' RR
MAFB
H3K27ac
(PC)
32
0
CTCF Motifs
Region
105,125,000
chr14
105,725,000
40,175,000
chr20
40,775,000

F

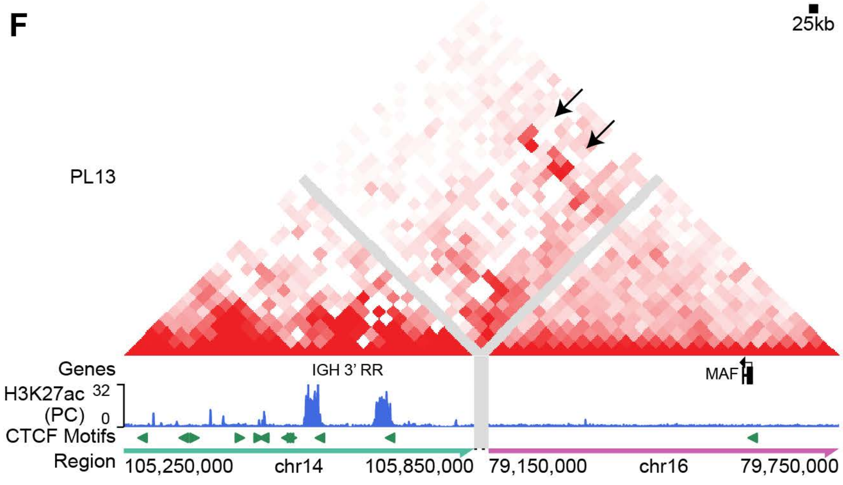
25kb
PL13
Genes
IGH 3' RR
MAF
H3K27ac
(PC)
32
0
CTCF Motifs
Region
105,250,000
chr14
105,850,000
79,150,000
chr16
79,750,000

G

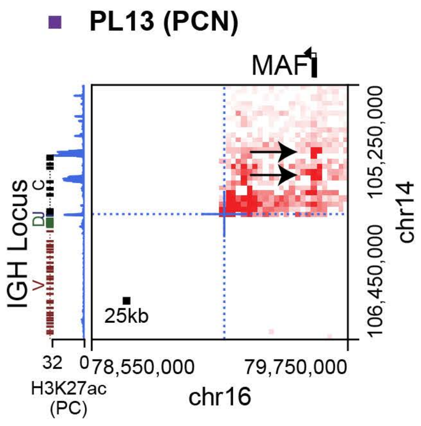
PL13 (PCN)
MAF
IGH Locus
C
DJ
V
25kb
105,250,000
106,450,000
chr14
32
0
78,550,000
79,750,000
H3K27ac
(PC)
chr16

#### Supplementary Figure S2

**A,** Balanced Hi-C contact matrices at 25kb resolution for all samples with *IGH::BCL2* rearrangements (corresponding with Fig 2G). Reference H3K27ac ChIP-Seq signal for germinal center B-cells (GCBC) is shown for the IGH locus.

**B,** Balanced Hi-C contact matrices at 25kb resolution for all samples with *IGH::CCND1* rearrangements (corresponding with Fig 2H). Reference H3K27ac ChIP-Seq signal for plasma cells (PC) is shown for the IGH locus.

**C,** Balanced Hi-C contact matrix at 25kb resolution, reconstructed across a chromosomal fusion between the *BCL2* and *IGL* loci in DL03 (corresponding with Figure 2I). Reference H3K27ac ChIP-Seq signal for GCBC is shown at bottom.

**D**, Balanced Hi-C contact matrix at 25kb resolution, reconstructed across a chromosomal fusion between the *IGH* and *CCND2* loci in PL08 (corresponding with Figure 2J). Reference H3K27ac ChIP-Seq signal for PC is shown at bottom.

**E**, Balanced Hi-C contact matrix at 25kb resolution, reconstructed across a chromosomal fusion between the *IGH* and *MAFB* loci in PL07 (corresponding with Figure 2K). Reference H3K27ac ChIP-Seq signal for PC is shown at bottom.

**F,** Balanced Hi-C contact matrix at 25kb resolution, reconstructed across a chromosomal fusion between the *IGH and MAF* loci in PL13. Reference H3K27ac ChIP-Seq signal for PC is shown at bottom.

**G,** Non-reconstructed balanced Hi-C contact matrix at 25kb resolution showing the *IGH::MAF* rearrangement in PL13. Reference H3K27ac ChIP-Seq signal for PC is shown for the *IGH* locus.

**Supplementary Figure S3**

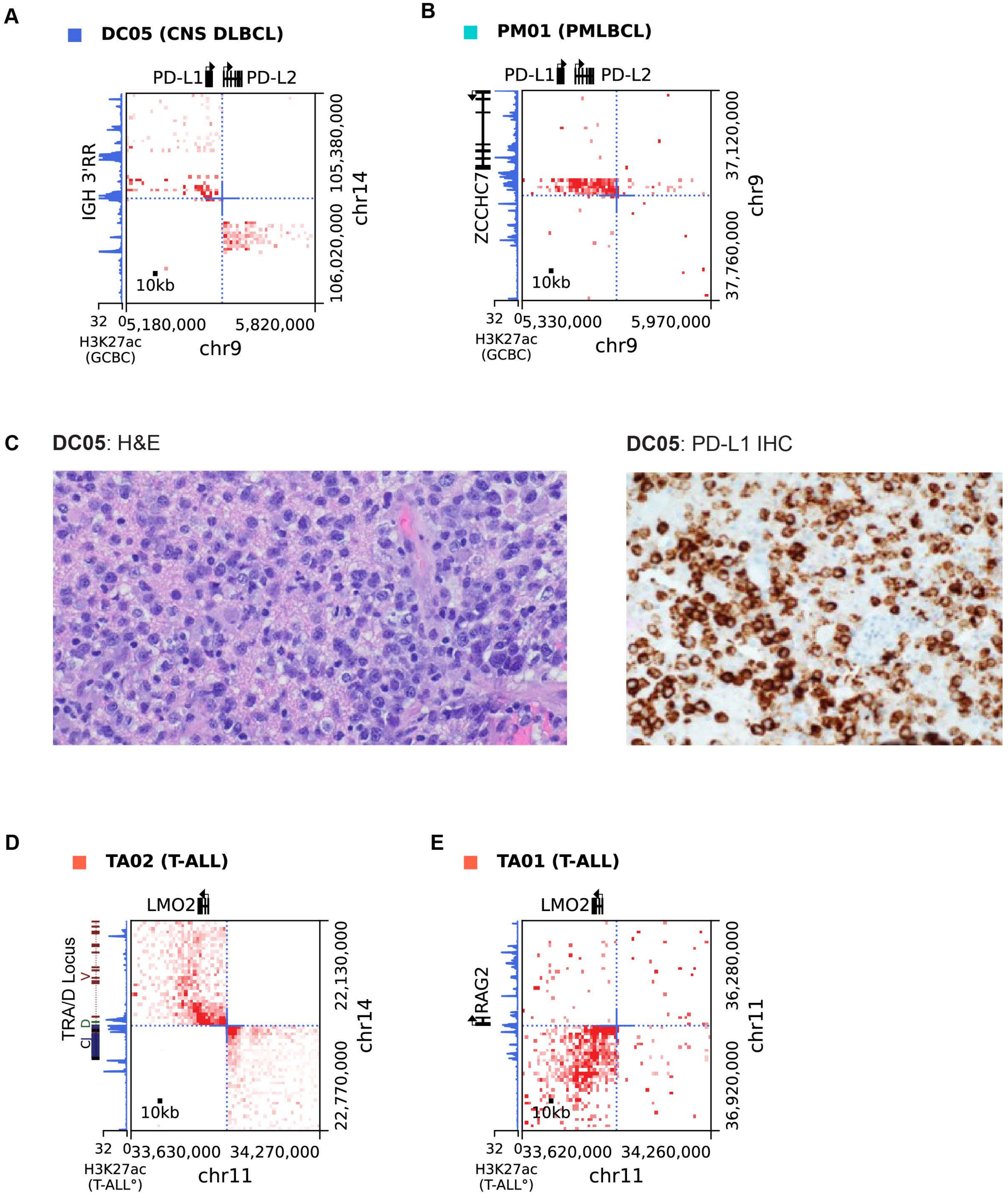

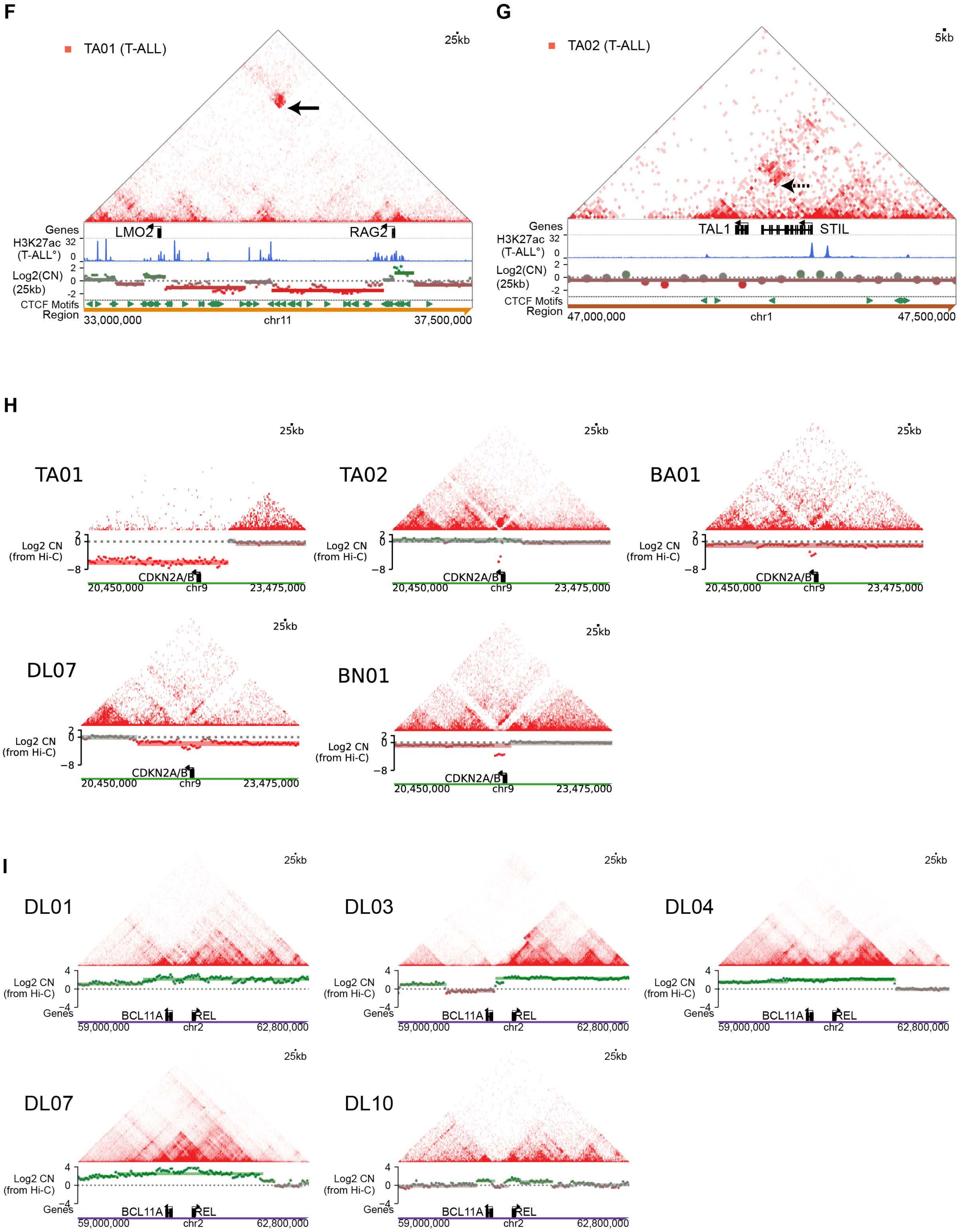
F
TA01 (T-ALL)
25kb
Genes
LMO2
RAG2
H3K27ac (T-ALL°)
32
0
Log2(CN) (25kb)
2
0
-2
CTCF Motifs
Region
33,000,000
chr11
37,500,000
G
TA02 (T-ALL)
5kb
TAL1
STIL
47,000,000
chr1
47,500,000
H
TA01
TA02
BA01
DL07
BN01
Log2 CN (from Hi-C)
-8
CDKN2A/B
20,450,000
chr9
23,475,000
I
DL01
DL03
DL04
DL07
DL10
4
-4
BCL11A
REL
59,000,000
chr2
62,800,000

#### ***Supplementary Figure S3***

**A**, Balanced Hi-C matrix at 10kb resolution showing chromosomal fusions between the *IGH* and *CD274* (PD-L1) loci in primary CNS large B cell lymphoma DC05.

**B**, Balanced Hi-C matrix at 10kb resolution showing chromosomal fusions between the *PAX5/ZCCHC7* and *PDCD1LG2* (PD-L2) loci in primary mediastinal large B-cell lymphoma PM01.

**C**, Photomicrograph of H&E and PD-L1 immunohistochemistry (positive) in DC05.

**D**, **E,** Balanced Hi-C matrices at 10kb resolution showing chromosomal fusions between the *LMO2* and *TRA/D* locus in TA02 (D), or LMO2 and *RAG2* loci in TA01 (E). Reference H3K27ac ChIP-Seq signal from a primary T-ALL sample is shown for the *TRA/D* and *RAG2* loci (from [98]).

**F** Raw Hi-C matrix for TA01 at 25kb depicting Hi-C signal corresponding with an *LMO2::RAG2* rearrangement. The solid arrow points to the aberrantly increased Hi-C signal between the fused regions that was called as a breakpoint by the automated callers. Hi-C-derived copy number profiles (calculated at 25kb resolution) are shown as tracks below (per-bin profiles as points, predicted segments as horizontal bars), with red corresponding to regions of copy number loss.

**G**, Raw Hi-C matrix for TA02 at 5kb resolution depicting Hi-C signal corresponding with a *TAL1::STIL* fusion. The dashed arrow points to the aberrantly increased Hi-C signal between the fused regions that was identified from manual review. Hi-C-derived copy number profiles (calculated at 25kb resolution) are shown as tracks below (per-bin profiles as points, predicted segments as horizontal bars), with red corresponding to regions of copy number loss.

**H,** Raw Hi-C matrices for samples with copy number loss of *CDKN2A/CDKN2B* by Hi-C. Copy number tracks are shown with green corresponding with log2 copy number above 0, and red corresponding with log2 copy number below 0, with color saturation increasing with greater absolute log2 copy number values.

**I**, Raw Hi-C matrices for samples with copy number gain of *REL* by Hi-C. Copy number tracks are shown with green corresponding with log2 copy number above 0, and red corresponding with log2 copy number below 0, with color saturation increasing with greater absolute log2 copy number values.

#### Supplementary Figure S4

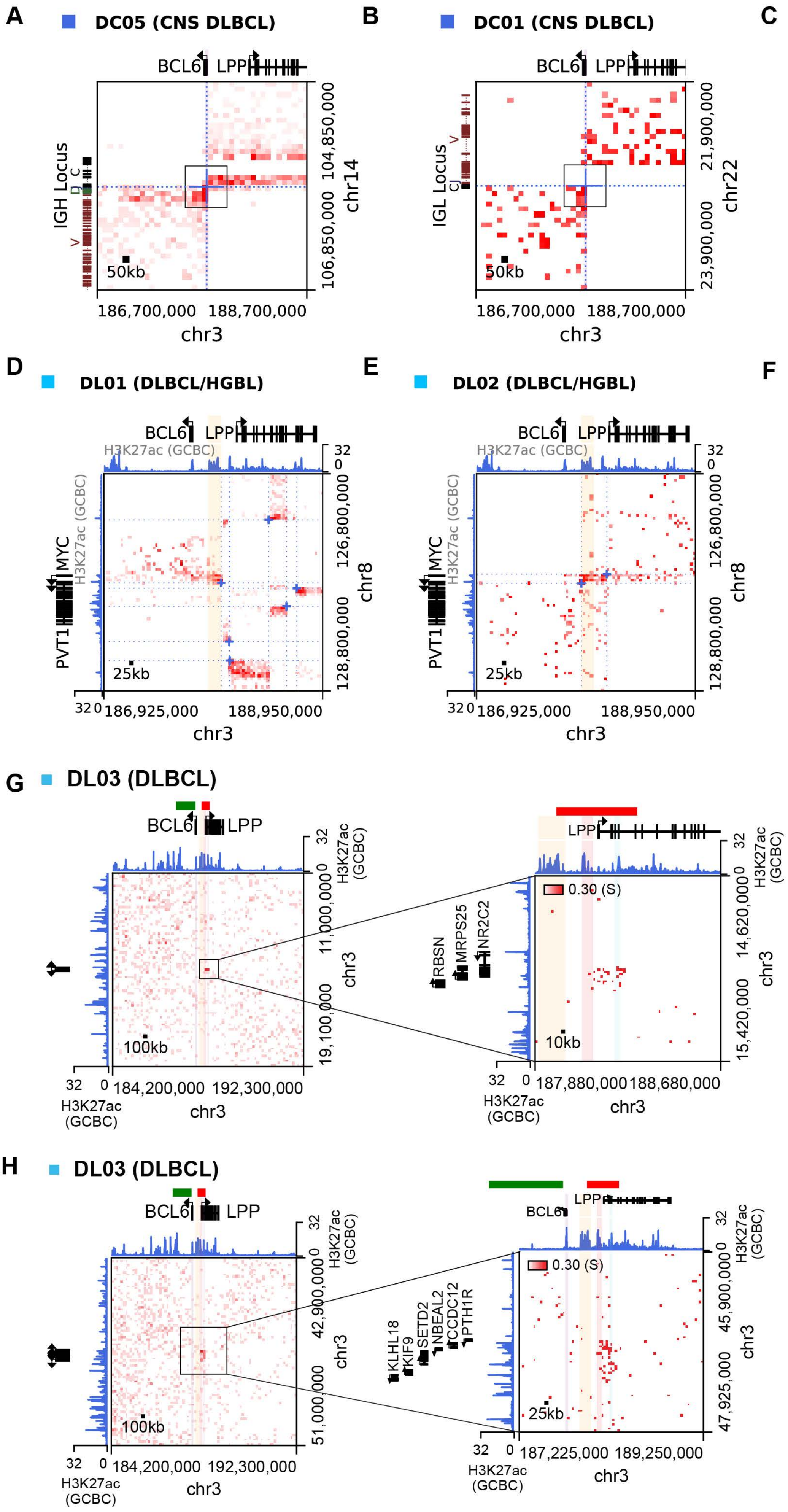

#### Supplementary Figure S4

**A, B, C**, Balanced Hi-C matrices at 50kb resolution showing rearrangements with the *BCL6* promoter region with the *IGH* locus*, IGL* locus and *JCHAIN* respectively. The boxed region corresponds with the zoomed Hi-C window used to visualize Figure 4B, 4C and 4D respectively.

**D**, **E,** Balanced Hi-C matrices at 25kb resolution showing *MYC::BCL6* super-enhancer rearrangements; the orange shaded region corresponds with the orange shaded region in Figure 4A.

**F**, Balanced Hi-C matrix at 25kb resolution depicting a *BCL6::BCL11A/REL* rearrangement.

**G, H**, Hi-C matrices showing possible insertion of a small genomic fragment containing the *LPP* promoter into two distant partner loci on chr3. The position of Vysis BCL6 break-apart FISH probes are shown in orange and green and reference GCB cell H3K27ac signal is shown in blue. Highlights show positions of the *BCL6* promoter (purple), *BCL6*-LCR super-enhancer (yellow) and two additional *BCL6* super-enhancer regions (pink and cyan). Note lack of heterologous interactions between either partner locus and the *BCL6* gene.

### Supplementary Figure S5

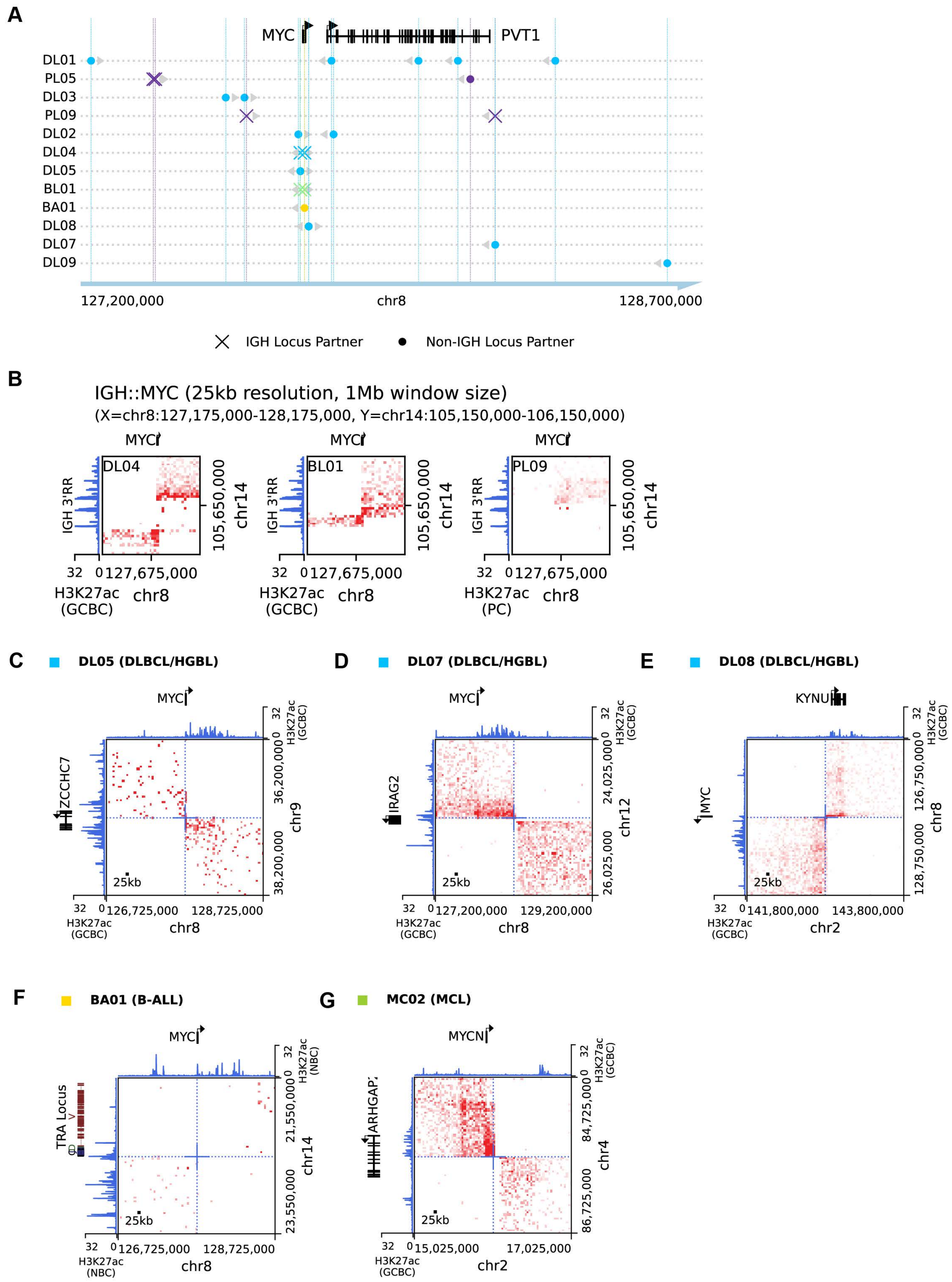

H

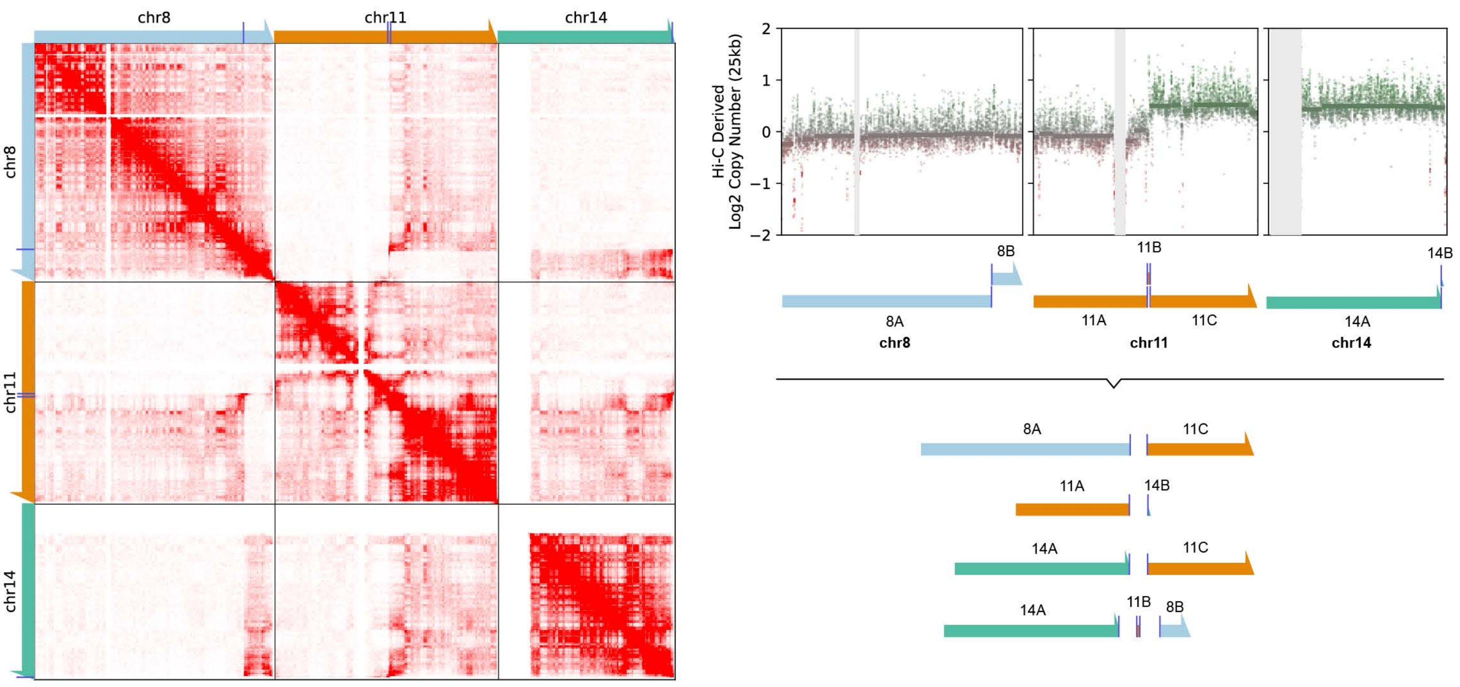

chr8
chr11
chr14
Hi-C Derived
Log2 Copy Number (25kb)
8A
8B
11A
11B
11C
14A
14B
chr8
chr11
chr14


I

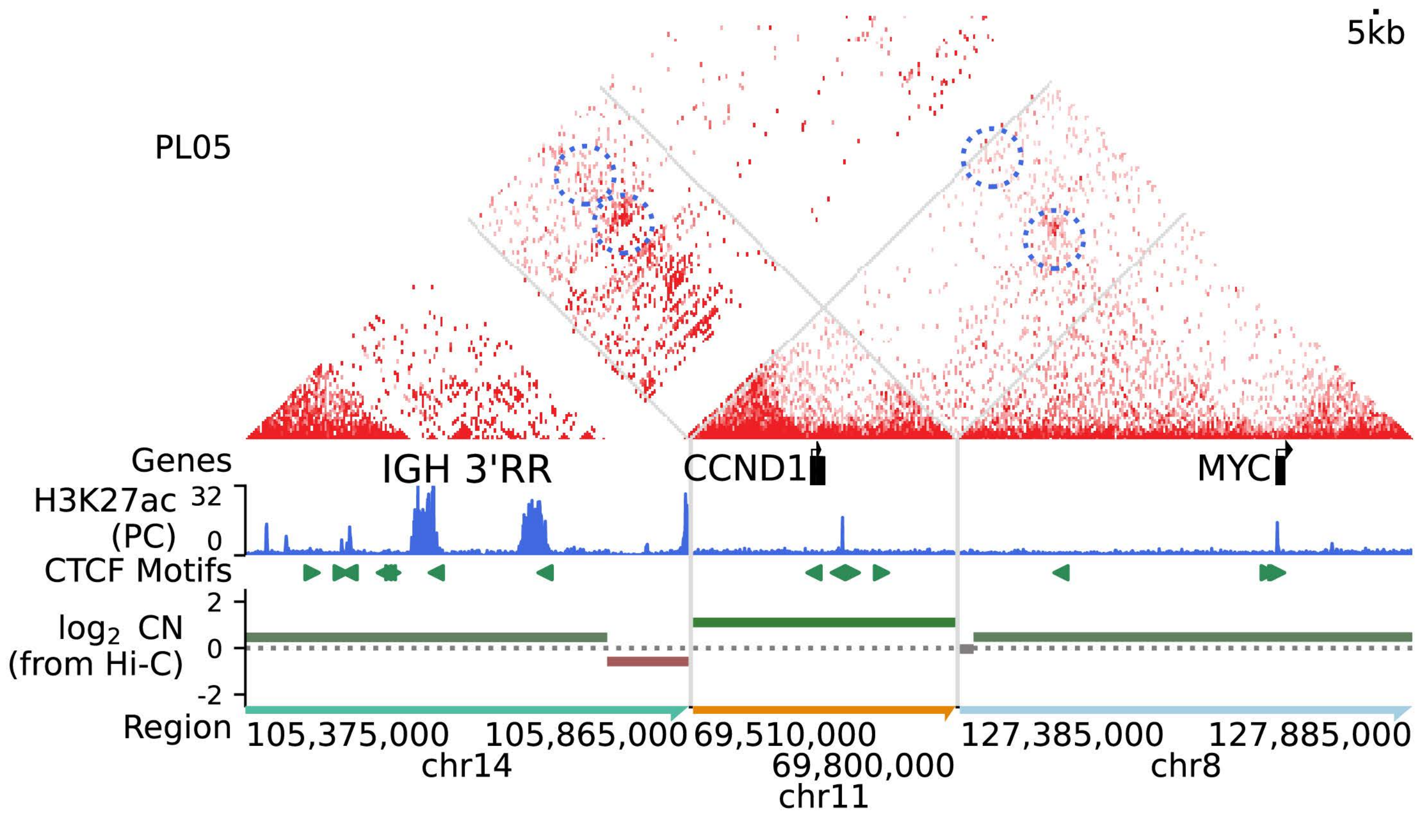

5kb
PL05
Genes
IGH 3'RR
CCND1
MYC
H3K27ac
(PC)
32
0
CTCF Motifs
log2 CN
(from Hi-C)
2
0
-2
Region
105,375,000
105,865,000
69,510,000
69,800,000
127,385,000
127,885,000
chr14
chr11
chr8

**J** **BA01** MYC Break-Apart FISH

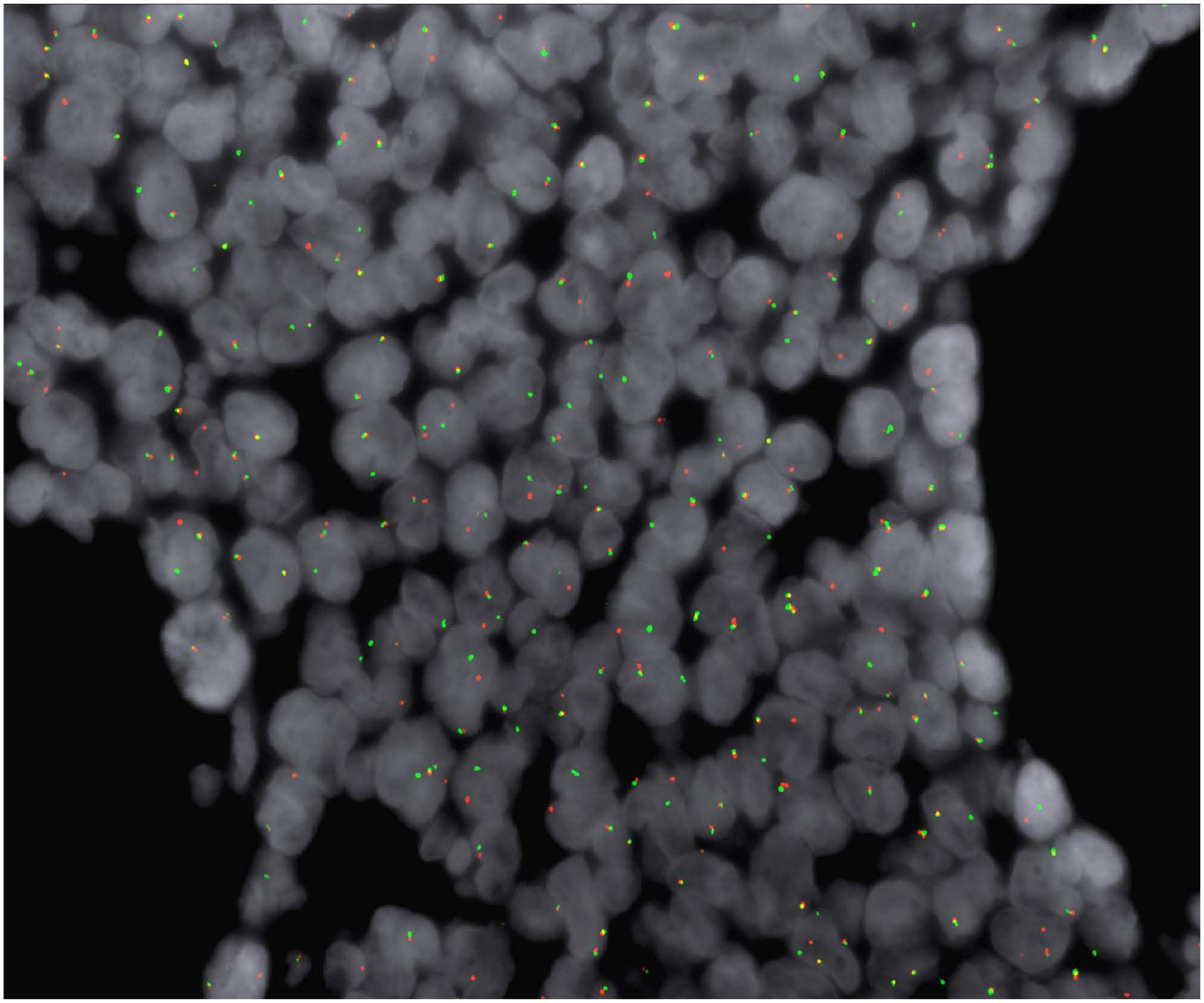

#### Supplementary Figure S5

**A**, Plot showing positions of all *MYC* locus rearrangement breakpoints relative to the *MYC* gene. Xs indicate *IGH* locus partners while circles indicate non-*IGH* locus partners. Grey arrowheads show breakpoint strandness (the direction/s of the genomic segment involved in the rearrangement/s). Marker colors denote diagnostic groups and follow the same scheme as Figure 1A and Supplementary Figure 1A.

**B**, Balanced Hi-C matrices at 25kb resolution showing simple *IGH::MYC* rearrangements in the indicated samples.

**C-E,** Balanced Hi-C matrices at 25kb resolution showing *MYC* rearrangements with non-*IGH* partner loci (*IRAG2, KYNU, PAX5/ZCCHC7*) to regions with active enhancers (represented by reference H3K27ac data). Panels C, D, and E correspond with Figures 5D, 5E and 5F respectively.

**F,** Balanced Hi-C matrix at 25kb resolution showing a *MYC* rearrangement with the *TRA* locus in a B-ALL sample that was identified on manual review. The Hi-C signal produced from this rearrangement is faint as the rearrangement is subclonal (see FISH in Supplementary Figure S5J).

**G,** Balanced Hi-C matrix at 25kb resolution showing a rearrangement between *MYCN* and *ARHGAP24* (corresponding with the Hi-C triangle in Figure 5G).

**H**, Data supporting reconstruction of 3-locus rearrangement between the MYC, IGH, and CCND1 loci in PCN biopsy PL05. Left, chr8, chr11 and chr14 from sample PL05 depicting raw Hi-C signal at 1Mb resolution for each chromosomal interaction and the location of breakpoints on chromosome schematics on each axis. Right, Hi-C derived copy number plot of chr8, chr11 and chr14 (25kb resolution), with schematic of chromosomal breakpoints and a possible reconstructed set of derivative chromosomes based on Hi-C interactions and large-scale copy number changes.

**I**, Balanced Hi-C data at 5kb resolution showing a possible reconstruction of the 3-way rearrangement involving *IGH, MYC* and *CCND1* in a PCN sample (PL05) with corresponding Hi-C derived copy number. Neo-loops are circled.

**J**, Composite fluorescence photomicrograph showing *MYC* break-apart FISH signals in a region of sample BA01. Note positive break-apart signals in a subset of nuclei (estimated at 15% throughout the biopsy)

#### Supplementary Figure S6

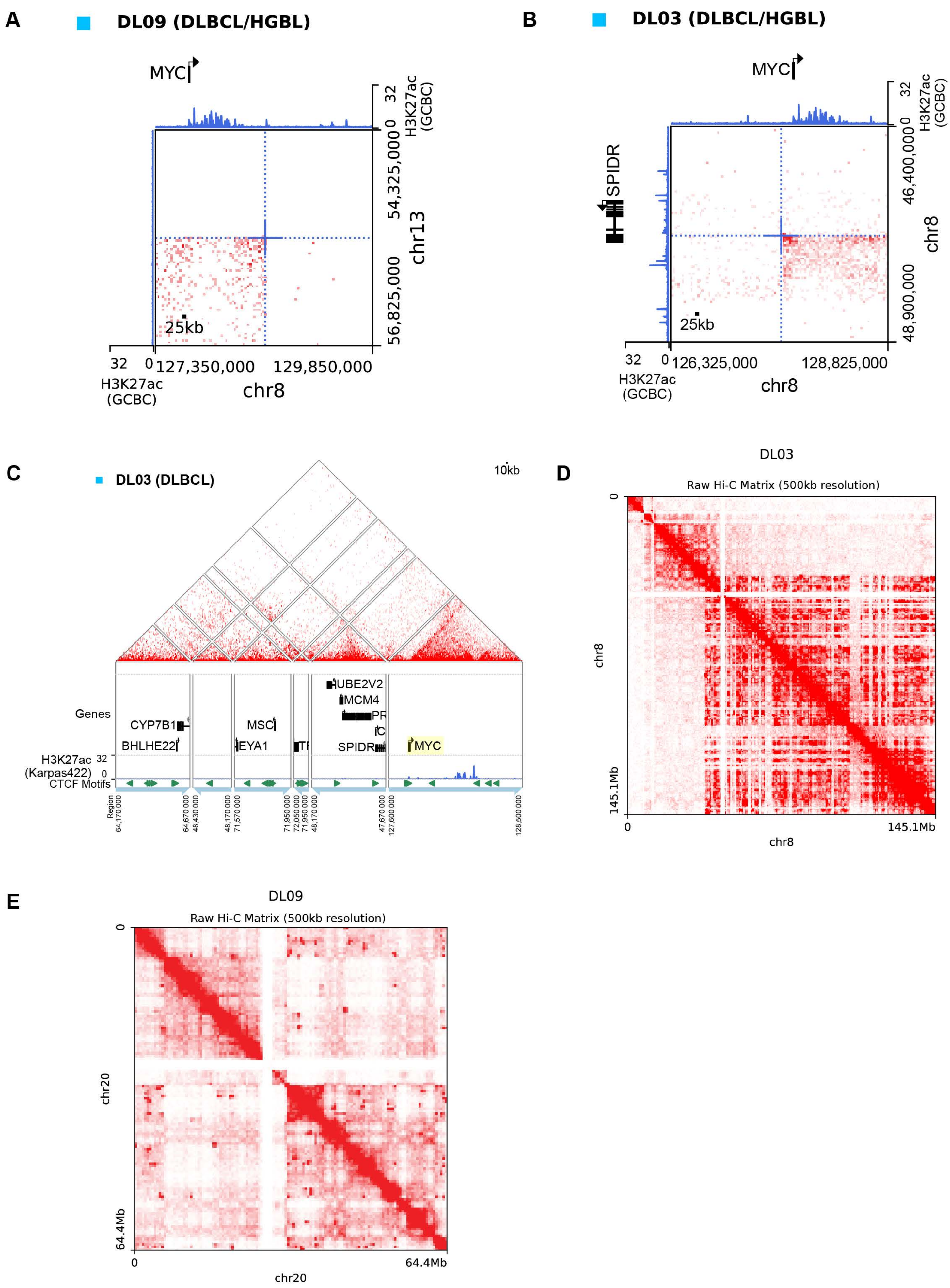

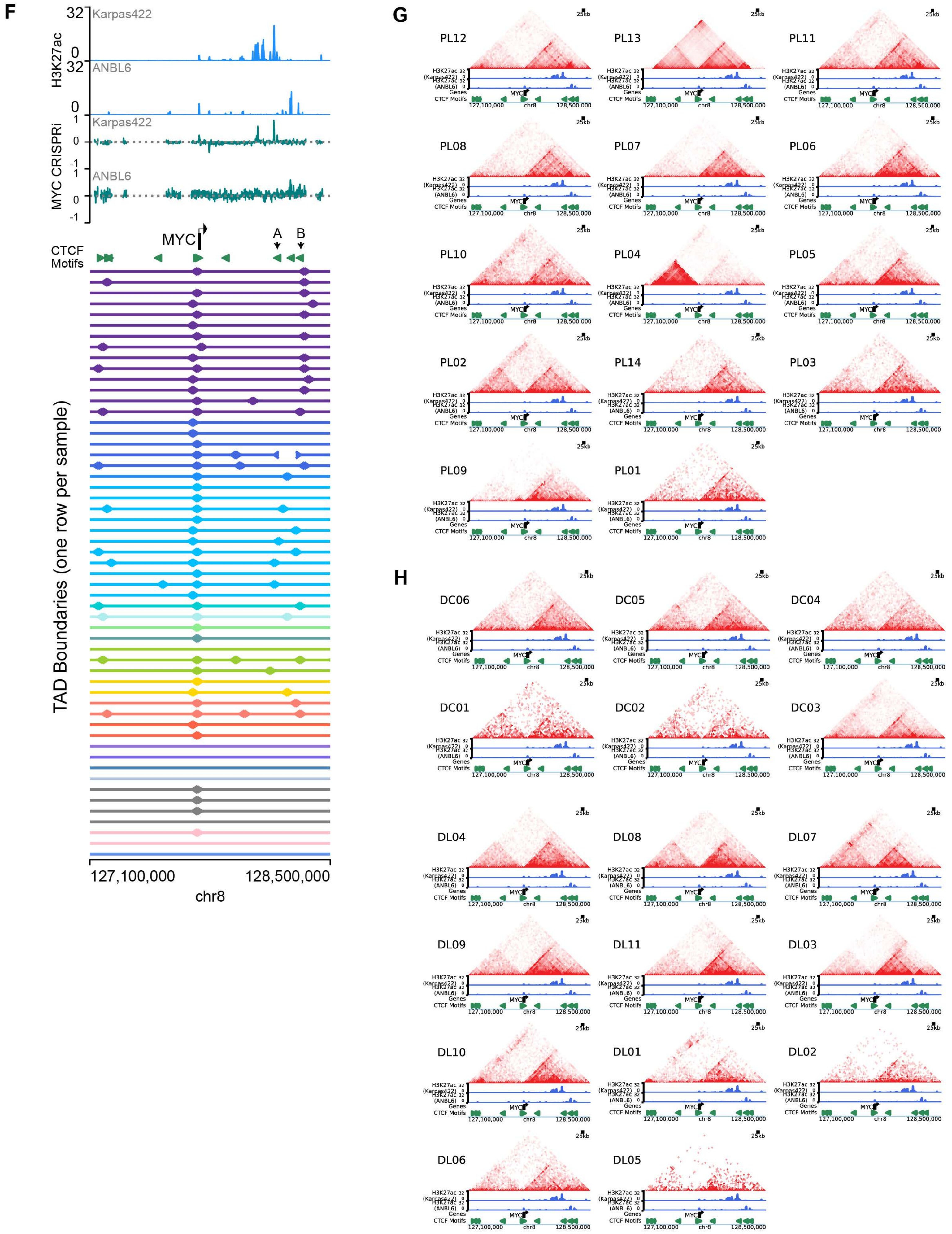
F
H3K27ac
Karpas422
ANBL6
MYC CRISPRi
Karpas422
ANBL6
MYC
A
B
CTCF Motifs
TAD Boundaries (one row per sample)
127,100,000
128,500,000
chr8
G
PL12
PL13
PL11
PL08
PL07
PL06
PL10
PL04
PL05
PL02
PL14
PL03
PL09
PL01
H
DC06
DC05
DC04
DC01
DC02
DC03
DL04
DL08
DL07
DL09
DL11
DL03
DL10
DL01
DL02
DL06
DL05
25kb
H3K27ac (Karpas422)
H3K27ac (ANBL6)
Genes
CTCF Motifs

#### Supplementary Figure S6

**A**, Balanced Hi-C matrix at 25kb resolution showing rearrangement between the *MYC* locus and a gene desert region of chromosome 13 in DL09, corresponding with Fig 6A.

**B**, Balanced Hi-C matrix at 25kb resolution showing an intrachromosomal rearrangement between the *MYC* and *SPIDR* loci, corresponding with Fig 6B.

**C**, Possible partial reconstruction of the derivative chr8 adjacent to the rearranged *MYC* locus based on intrachromosomal breakpoints in DL03. The Hi-C signal at each breakpoint is traced backwards and sudden loss of Hi-C interaction signal which matches the position of aberrant gain of Hi-C interaction signal at another breakpoint indicates a point of fusion. Breakpoint anchors are followed backwards successively in a chain to produce a putative reconstruction.

**D**, Raw Hi-C matrix at 500kb resolution showing chromothripsis of chr8 in DL03.

**E**, Raw Hi-C matrix at 500kb resolution showing chromothripsis of chr20 in DL09.

**F,** Position of state-specific MYC enhancers with regard to FFPE Hi-C TAD boundaries in in the genomic region chr8:127,100,000-128,500,000 (corresponding with Figure 6C). Top: H3K27ac ChIP-Seq signal and tiling CRISPRi screen score (-log2 depletion, 20 sgRNA sliding window) for the GCB-DLBCL cell line Karpas-422 and the MM cell line ANBL6. Bottom: TAD boundaries at 25kb resolution in FFPE Hi-C datasets ordered and colored as in Figure 1A.

**G, H**, Individual raw Hi-C contact matrices at 25kb resolution across the MYC locus in all PCN samples (G) and CNS and non-CNS DLBCL samples (H), corresponding with the same region as Figure 6C. Reference H3K27ac data is shown for Karpas422 and ANBL6 cell lines as well as oriented CTCF motifs. Samples are ordered as in panel F.
